# The Arabidopsis condensin CAP-D subunits arrange interphase chromatin

**DOI:** 10.1101/2019.12.12.873885

**Authors:** Celia Municio, Wojciech Antosz, Klaus D. Grasser, Etienne Kornobis, Michiel Van Bel, Ignacio Eguinoa, Frederik Coppens, Andrea Bräutigam, Inna Lermontova, Astrid Bruckmann, Andreas Houben, Veit Schubert

## Abstract

Condensins are best known for their role in shaping chromosomes. However, other functions as organizing interphase chromatin and transcriptional control have been reported in yeasts and animals. Yeasts encode one condensin complex, while higher eukaryotes have two of them (condensin I and II). Both, condensin I and II, are conserved in *Arabidopsis thaliana*, but so far little is known about their function. Here we show that the *A. thaliana* CAP-D2 (condensin I) and CAP-D3 (condensin II) subunits are highly expressed in mitotically active tissues. *In silico* and pull-down experiments indicate that both CAP-D proteins interact with the other condensin I and II subunits. Our data suggest that the expression, localization and composition of the condensin complexes in *A. thaliana* are similar as in other higher eukaryotes. Previous experiments showed that the lack of *A. thaliana* CAP-D3 leads to centromere association during interphase. To study the function of CAP-D3 in chromatin organization more in detail we compared the nuclear distribution of rDNA, of centromeric chromocenters and of different epigenetic marks, as well as the nuclear size between wild-type and *cap-d3* mutants. In these mutants an association of heterochromatic sequences occurs, but nuclear size and the general methylation and acetylation patterns remain unchanged. In addition, transcriptome analyses revealed a moderate influence of CAP-D3 on general transcription, but a stronger one on transcription of stress-related genes. We propose a model for the CAP-D3 function during interphase, where CAP-D3 localizes in euchromatin loops to stiff them, and consequently separates centromeric regions and 45S rDNA repeats.

## INTRODUCTION

The spatial genome arrangement is important to regulate the access of proteins to DNA (Gibcus and Dekker, 2013), since the folding of chromatin allows or impedes interactions between distinct loci and their regulatory sequences (Doğan and Liu, 2018; Robson *et al.*, 2019; Stam *et al.*, 2019; Szabo *et al.*, 2019). Thus, a better knowledge of the nuclear organization during interphase could help to understand processes like replication, DNA repair, recombination and transcription. During interphase, in *A. thaliana* (Pecinka *et al.*, 2004) as well as in other higher eukaryotes, the chromosomes occupy discrete regions called chromosome territories. Although this higher order chromatin arrangements is highly conserved, species-specific structural and functional features of nuclear organization exist in metazoans and protists (Cremer and Cremer, 2010; Cremer *et al.*, 2018). In contrast to mammals (Boyle *et al.*, 2001; Mayer *et al.*, 2005) and birds (Habermann *et al.*, 2001), *A. thaliana* chromosome territories prefer no particular position within the nucleus (Pecinka *et al.*, 2004).

Chromocenters are chromatin structures intensely stained by DNA-specific dyes and represent condensed heterochromatin regions in interphase nuclei (Jost *et al.*, 2012). In *A. thaliana* chromocenters incarnate centromeric and pericentromeric heterochromatin located near the nuclear periphery and the nucleolus (Fransz *et al.*, 2002; Schubert *et al.*, 2012). For the maintenance of chromocenters different proteins related to methylation, ATPases and nuclear periphery components were described in *A. thaliana* (Soppe *et al.*, 2002; Moissiard *et al.*, 2012; Wang *et al.*, 2013; Poulet *et al.*, 2017).

To explain the organization of chromosome territories in *A. thaliana* interphase nuclei, a rosette model was proposed (Fransz *et al.*, 2002). Based on cytological observation and later support by computer simulations (de Nooijer *et al.*, 2009) and Hi-C data (Feng *et al.*, 2014; Liu *et al.*, 2016), this model assumes that the chromosomes are organized as chromatin loops emanating from the chromocenters. Structural Maintenance of Chromosomes (SMC) complexes are present in prokaryotes and eukaryotes (Cobbe and Heck, 2004). They are essential for chromatin organization and dynamics, gene regulation and DNA repair. In eukaryotes six conserved SMC subunits form the core of three different complexes: cohesin, involved in sister chromatids cohesion and interphase chromatin arrangement; condensin, involved in mitotic and meiotic chromosome organization (van Ruiten and Rowland, 2018; Skibbens, 2019); and the SMC5/SMC6 complex, mainly involved in DNA repair and replication (Jeppsson *et al.*, 2014). Animals have two condensin complexes, condensin I and II (Ono *et al.*, 2003). In yeasts, only one condensin complex analogous to animal condensin I is present (Freeman *et al.*, 2000; Hirano, 2012a). Condensin I and II share a core formed by SMC2 and SMC4 and differ in the associated proteins, which are in condensin I CAP-H, CAP-D2 and CAP-G, and in condensin II CAP-H2, CAP-D3 and CAP-G2 (Ono *et al.*, 2003; Hirano, 2012a). This composition is conserved in higher eukaryotes, although in Drosophila the subunit CAP-G2 of condensin II has not been detected (Herzog *et al.*, 2013). As proposed for *A. thaliana* (Fig. 1), plants apparently have condensin I and II.

**Figure 1.**
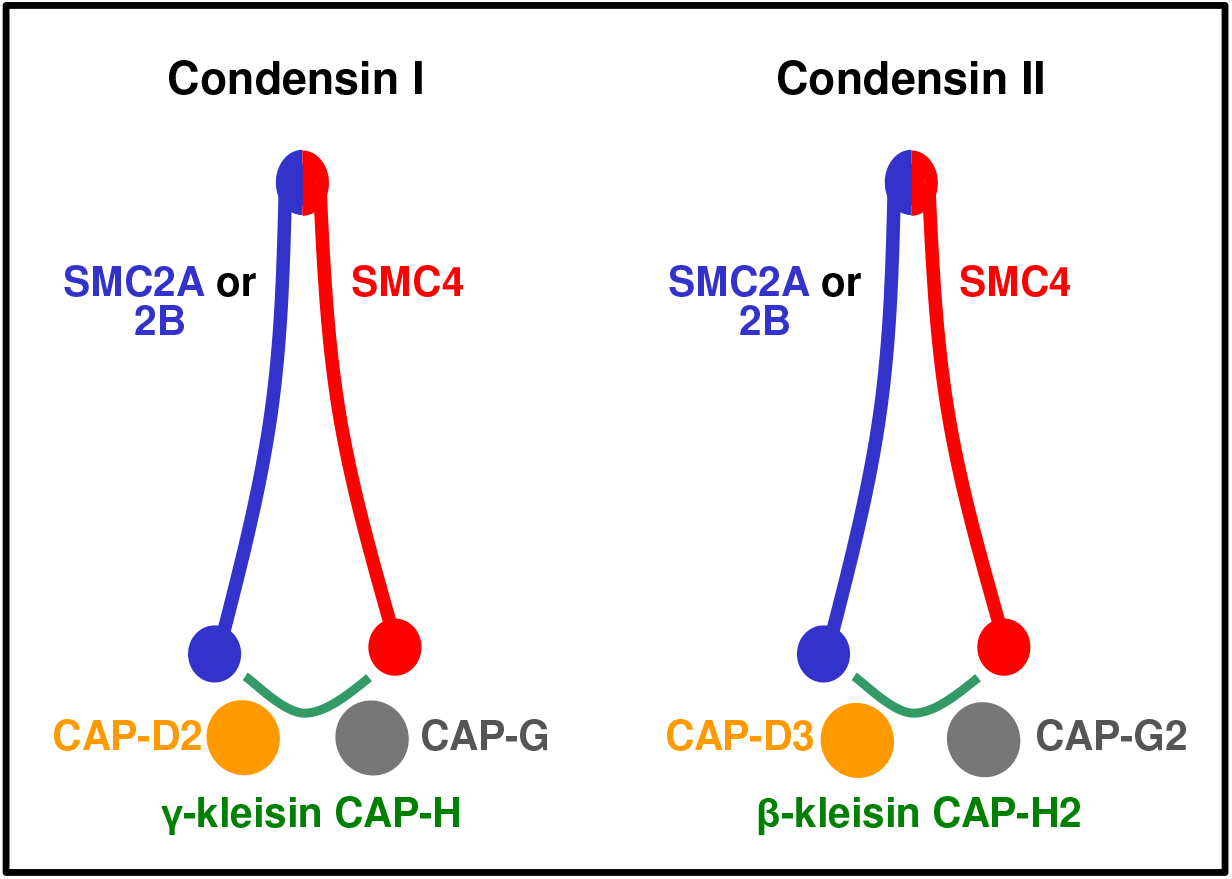
*A. thaliana* condensin I and II subunit composition based on models of Nasmyth and Hearing (2005) and Schubert (2009). Both condensin complexes can be formed presumably by SMC4 and two alternative SMC2 subunits. Condensin I contains in addition CAP-D2, CAP-G and the γ-kleisin CAP-H, condensin II CAP-D3, CAP-G2 and the β-kleisin CAP-H2 (www.arabidopsis.org; Fujimoto et al., 2005). In the present work we confirm via analyzing CAP-D2 and CAP-D3 the interaction with the other respective subunits, and thus the presence of condensin I and II in *A. thaliana*.

Condensins have been widely studied in human, animals and yeast for their role in shaping chromosomes. Together with topoisomerase II condensins form a scaffold within human somatic metaphase chromatids (Maeshima and Laemmli, 2003). Depletion of condensin I causes short fuzzy metaphase chromosomes while the depletion of condensin II causes long and curly chromosomes (Ono *et al.*, 2003; Green *et al.*, 2012). Besides aberrant chromosome morphologies, chromosomes lacking several condensin subunits show anaphase bridges and other segregation defects (Freeman *et al.*, 2000; Hudson *et al.*, 2003; Ono *et al.*, 2003, 2004; Hirota *et al.*, 2004; Savvidou *et al.*, 2005; Gerlich *et al.*, 2006; Hartl *et al.*, 2008). Both complexes may form DNA loops resulting in chromosome compaction (Elbatsh *et al.*, 2019; Gibcus *et al.*, 2018; van Ruiten and Rowland, 2018; Walther *et al.*, 2018).

Condensin I and II complexes show a distinct subcellular localization during the mammalian cell cycle. In human and rat during interphase, condensin I occurs in the cytoplasm, condensin II in the nucleus (Hirota *et al.*, 2004; Ono *et al.*, 2004). During mitosis, condensin I and II localize along the chromosome arms in an alternate fashion, and both are enriched at the centromeres (Ono *et al.*, 2003; Ono *et al.*, 2004; Savvidou *et al.*, 2005). In addition to the canonical role in metaphase chromosome formation, condensins are also involved in gene expression and chromatin organization during interphase (Wallace and Bosco, 2013; Wallace *et al.*, 2015). In mouse and human, condensin II localizes at the promoters of active genes and is required for normal gene expression (Dowen *et al.*, 2013; Yuen *et al.*, 2017;). In *Drosophila*, CAP-D3 together with the RetinoBlastoma protein RBF1, regulates gene clusters involved in tissue-specific functions (Longworth *et al.*, 2012), and condensin II promotes the formation of chromosome territories and keeps repetitive sequence clusters apart from each other (Hartl *et al.*, 2008; Bauer *et al.*, 2012; Hirano, 2012b; Rosin *et al.*, 2018).

Also *A. thaliana* posseses the components for both condensin complexes (Schubert, 2009; Smith *et al.*, 2014). In contrast to other organisms, *A. thaliana* has two SMC2 homologs, SMC2A and SMC2B with redundant functions (Siddiqui *et al.*, 2003)(Fig. 1). As in other species, SMC4, CAP-H and CAP-H2 are present within chromosomes and are required for normal metaphase chromosome compaction (Fujimoto *et al.*, 2005; Smith *et al.*, 2014). During interphase, CAP-H is present in the cytoplasm of protoplasts while the condensin II subunits CAP-H2 and CAP-D3 were detected in the nucleolus and euchromatin, respectively (Fujimoto *et al.*, 2005; Schubert *et al.*, 2013). CAP-D2 and CAP-D3 prevent the association of centromeres and induce chromatin compaction (Schubert *et al.*, 2013). The requirement of the condensin II-specific subunits CAP-H2 and CAP-G2 for keeping centromeres apart has been confirmed by Sakamoto *et al.* (2019). In addition, these authors showed that condensin II is necessary for the correct spatial arrangement between centromeres and rDNA arrays.

Condensins are highly conserved, but have not been studied extensively in plants. Here we analyze the *A. thaliana* CAP-D2 and CAP-D3 condensin subunit expression patterns, their cellular localization and interaction with other condensin subunits and additional proteins for better understanding their functions. We demonstrate that also *A. thaliana* forms specific condensin I and II complexes, and show that CAP-D3 mediates the spatial separation of chromocenters, without altering the global methylation pattern and nuclear ultrastructure. Finally, we suggest a model explaining the action of CAP-D3 to prevent the association of chromocenters.

## RESULTS

### *CAP-D2* and *CAP-D3* are highly expressed in meristematic tissues

Based on *in silico* analysis using the Arabidopsis eFP Browser (Winter *et al.*, 2007), *A. thaliana CAP-D2* (At3g57060) and *CAP-D3* (At4g15890) have a similar expression pattern. Both proteins are highly expressed in the shoot apex, roots, flower buds and vegetative rosette leaves. Their expression is lower in cotyledons, rosette leaves after bolting, mature flowers, siliques and embryos (Figure S1). To corroborate the *in silico* data we assessed the transcription of both genes in seedlings, mature rosette leaves, roots and flower buds by quantitative real-time RT-PCR. The highest transcription of both genes was observed in flower buds, the lowest in seedlings. The transcription level of *CAP-D2* is 25.6, 14.8 and 3.5 times higher in flower buds, roots and leaves, respectively, than in seedlings. Similarly, the *CAP-D3* transcription is 18.3, 9.4 and 4.4 times higher in flower buds, roots and leaves respectively, than in seedlings (Fig. 2).

**Figure 2.**
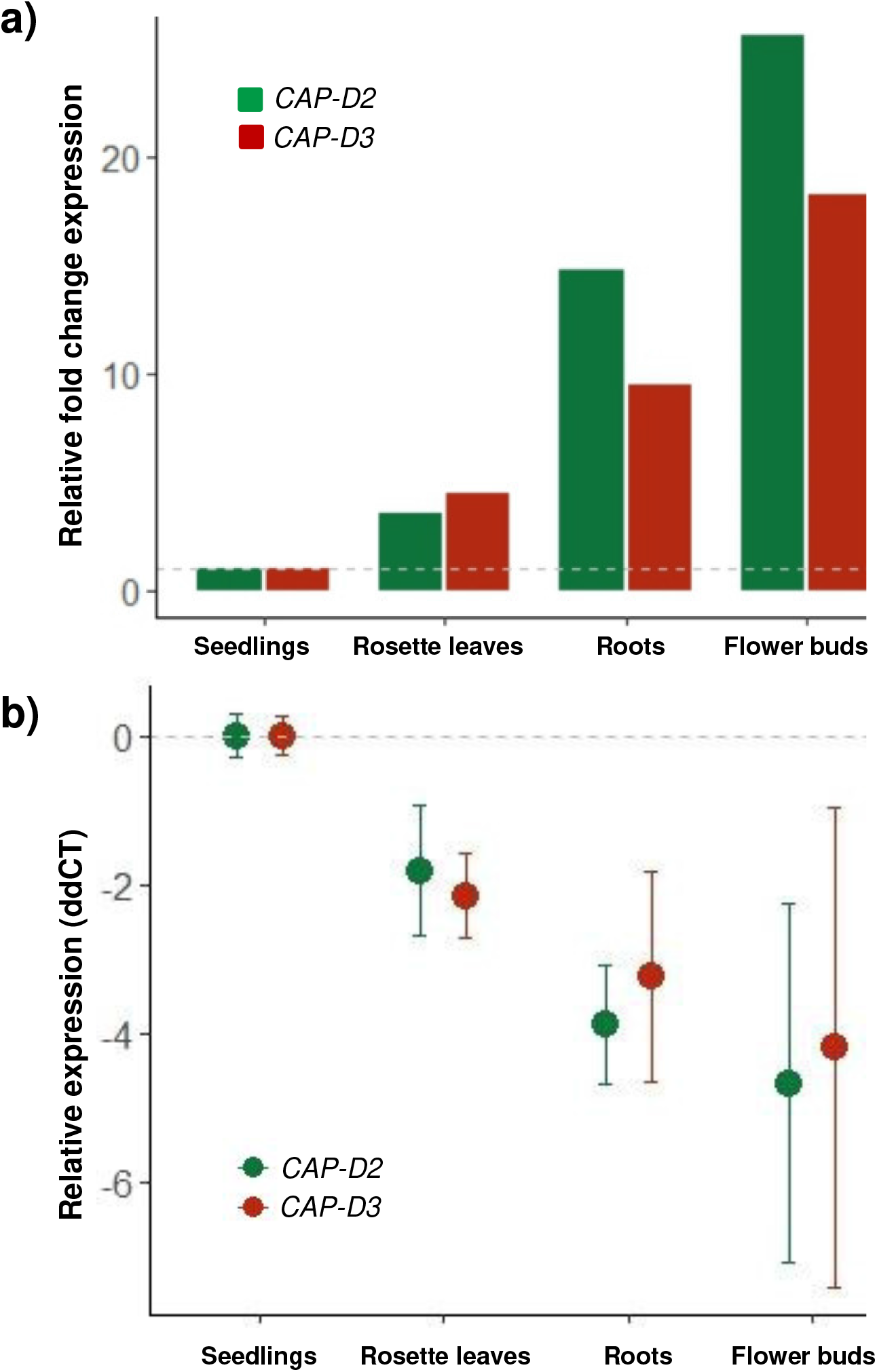
Transcription of *CAP-D2* and *CAP-D3* in different tissues. Relative fold change expression **(a)** and relative expression (ddCT) **(b)** in flower buds, roots and rosette leaves compared to seedlings. The values were normalized to the geometric mean of the house keeping genes *PP2A* and *RHIP1* and relative to the expression in seedlings. Lower ddCT values indicate higher transcription. Error bars in **(b)** represent the standard deviation between three biological replicates (each in triplicates). No error bars are shown in **(a)** since the fold change is a direct conversion of the ddCTvalues.

The activity of the *CAP-D2* and *CAP-D3* promoters was evaluated in *A. thaliana* transgenic lines expressing different versions of the promoters fused to the β-glucuronidase (GUS) reporter gene (Fig. 3a). Six presumed promoters of different length were analyzed for *CAP-D2* and two for *CAP-D3*. The promoter region of *CAP-D2* contains two putative E2F binding sites at −345 bp and −114 bp upstream from the start of the gene (Schubert *et al.*, 2013). Two promoter lengths were analyzed: a promoter that comprises 1156 bp upstream of the start of *CAP-D2* (Pro4), and a short promoter of 391 bp (Pro7). In addition, promoter proximal introns can enhance the expression of a gene by a mechanism known as Intron-Mediated Enhancement (IME) (Rose *et al.*, 2008). The putative enhancing ability of *CAP-D2* introns was analyzed *in silico* with the web tool IMEter (Parra *et al.*, 2011). The IMEter score is positively correlated to the enhancing ability of an intron. For *CAP-D2* the two first introns have positive IMEter scores of 12.13 and 2.36, respectively. Thus, it is likely that they enhance expression. These introns were included in the analysis in combination with the long and short promoters of *CAP-D2*: Pro5 (long promoter) and Pro8 (short promoter) include Intron1; and Pro6 (long promoter) and Pro9 (short promoter) include both Intron1 and Intron2. The promoter region of *CAP-D3* contains also two putative E2F binding sites at −397 bp and −84 bp (Schubert *et al.*, 2013). However, the IMEter scores of the first two *CAP-D3* introns were negative, −13.20 and −5.88 respectively. Thus, it is unlikely that they enhance expression. Therefore, for *CAP-D3* the introns were not considered and only a long promoter at −1318 bp (Pro10) and a short promoter at −474 bp (Pro11) from the start of the gene were analyzed.

**Figure 3.**
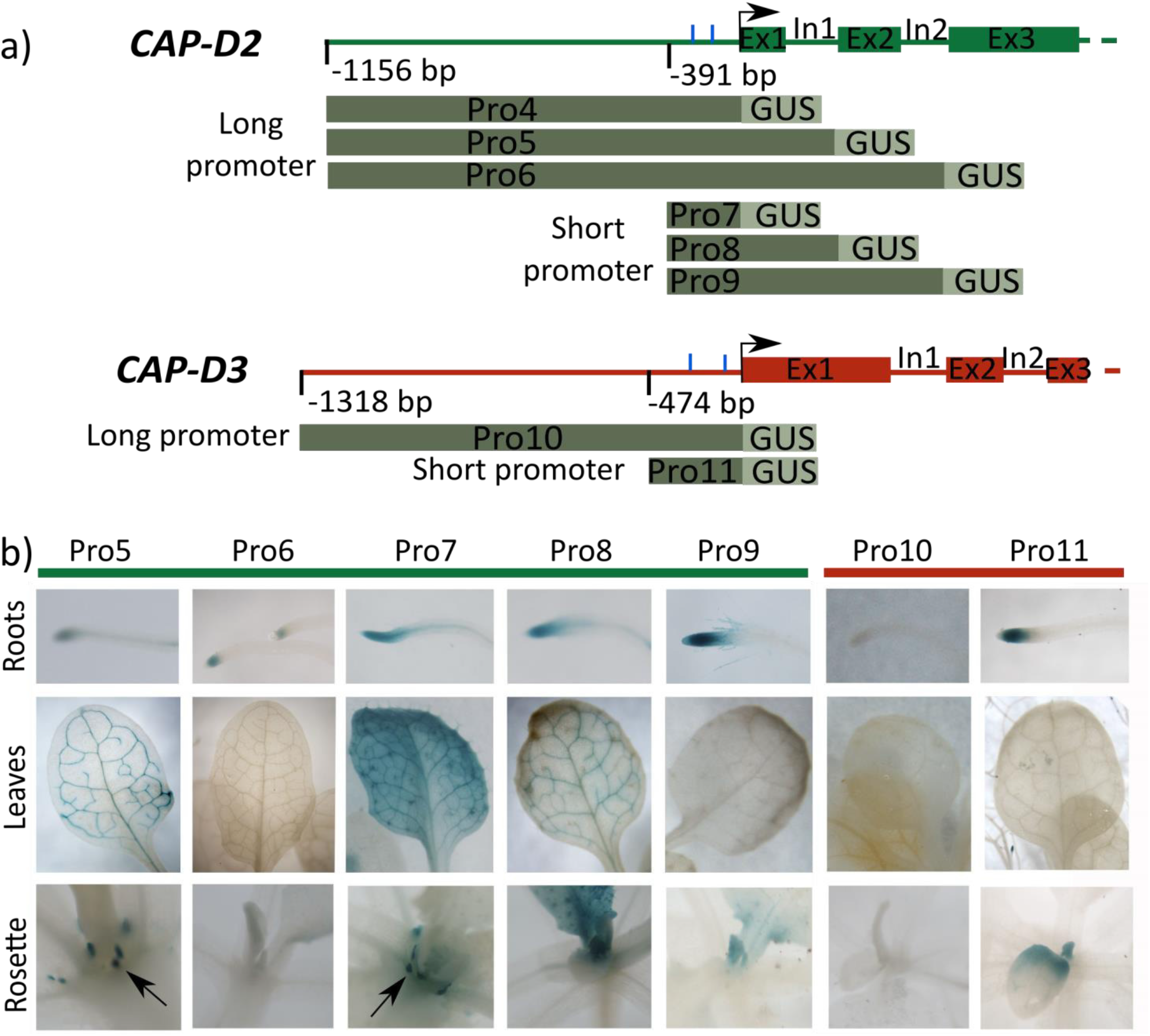
*CAP-D2* and *CAP-D3* promoter activity. **a)** Schemata of the promoter regions, the first two introns and three exons of *CAP-D2* and *CAP-D3*. The start of the coding region is marked by an arrow and the blue lines represent the position of the E2F binding sites. Below are shown the different tested promoter versions fused to the GUS gene. **b)** Histochemical GUS staining (blue staining) in root meristems, leaves, stipules (arrows; Pro5, 7) and apical meristems of plants transformed with the indicated promoter versions Pro5-9 and Pro10-11. Images of Pro4 are not depicted since no transformants could be isolated.

T1 transgenic plants with the different versions of *CAP-D2* and *CAP-D3* promoters were stained for GUS analysis (Fig. 3b). Only for Pro4 no positive plants could be isolated. The *CAP-D2* promoter version Pro5 (n=7) was active in stipules (small organs at the base of the leaves), leaf vascular tissue and root tip meristems. Pro6 (n=6) had weak activity in root tips. All Pro7 plants (n=21) showed GUS-staining in leaf vascular tissue and root tips, and 16 plants also in stipules. All Pro8 plants (n=23) presented GUS activity in the apical meristem and root tips, and 16 of them also in leaf vascular tissue. Pro9 (n=5) showed activity in roots, and 3 plants also weakly in the apical meristem (Fig. 1d). Therefore, all *CAP-D2* promoter versions were active in root tips, but the staining was stronger in the short promoter versions (Pro7, Pro8 and Pro9) than in the long ones (Pro5 and Pro6). In addition, the *CAP-D2* short promoters showed an activity in the apical meristem and versions that included the second intron (Pro6 and Pro9) lost the staining in the leaf vascular tissue. *CAP-D3* Pro10 showed no activity, and for Pro11 (n=8), the plants showed activity in the apical meristem and root tips. For both, *CAP-D2* and *CAP-D3*, the expression can be driven more effectively by the short promoter, which contains the E2F sites.

Taken together quantitative real-time RT-PCR and GUS activity staining demonstrated that, *CAP-D2* and *CAP-D3* are highly expressed in meristematic tissues (root tip meristem, flower buds, apical meristem) and young leaves and less expressed in mature leaves. The low transcription observed in seedlings could be due to a low amount of meristematic tissue in the sample since just one-week old seedlings were used for RNA isolation.

### CAP-D2 and CAP-D3 interact with the other condensin subunits in specific complexes

CAP-D2 and CAP-D3 are specific components of condensin I and II complexes, respectively. The presence of CAP-D2 and CAP-D3 as well as the other condensin complex subunits in *A. thaliana* was previously confirmed (Smith *et al.*, 2014), but whether the complexes are formed by the same subunits as in non-plant species is unknown. To predict a composition of each complex we identified putative interactors of CAP-D2 (Figure S2a) and CAP-D3 (Figure S2b) *in silico* using the STRING program (http://string-db.org/; Szklarczyk *et al.*, 2019). At the high score of >0.90 the following proteins: SMC2A (At5g62410), SMC2B (At3G47460) and SMC4 (At5g48600) were identified in interaction networks of both CAP-D2 and CAP-D3, while CAP-G (At5g37630) and CAP-H (At2g32590) were found as interactors of CAP-D2, and CAP-G2 (At1g64960) and CAP-H2 (At3g16730) as specific interactors of CAP-D3, respectively. Due to the presence of SMC2A, SMC2B and SMC4 in both interaction networks they may be involved in the formation of condensin I as well as of condensin II. *In silico* analysis using the STRING program identified besides cohesin subunits also SMC5/6 complex subunits as CAP-D2 and CAP-D3 interacting partners (Zelkowski *et al.*, 2019).

To confirm these *in silico* results and to determine the composition of each complex experimentally, CAP-D2 and CAP-D3 were fused to a GS-tag, and affinity-purified from *A. thaliana* PSB-D suspension cultured cells (Figure S3). The proteins co-purifiying with CAP-D2-GS and CAP-D3-GS were identified by mass spectrometry. The putative subunits of the condensin I complex, SMC2A, SMC2B, SMC4, CAP-H and CAP-G, were detected with high scores in the CAP-D2-GS eluates of three affinity purifications performed. Similarly, the putative subunits of the condensin II complex, SMC2A, SMC4, CAP-H2 and CAP-G2, were detected in the three affinity purifications performed for CAP-D3-GS and SMC2B in two of the affinity purifications (Table 1, Fig. 1). Like in the *in silico* analysis, CAP-H, CAP-G and CAP-H2, CAP-G2 were identified as specific components of the condensin I and condensin II complexes, respectively, while SMC2A, SMC2B and SMC4 co-precipitated with both CAP-D2 and CAP-D3.The results indicate that *A. thaliana*, similar as mammals, chicken and *C. elegans* (Hirano, 2012a; Onn *et al.*, 2007), comprises specific condensin I and II complexes. Interestingly, in addition to SMC4, both SMC2A and SMC2B may be involved in the formation of both condensin complexes.

**Table 1.** Condensin subunits co-purifying with CAP-D2 and CAP-D3.

| Bait | Interactor (AGI) | Times detected | Average MASCOT Score |
| --- | --- | --- | --- |
| CAP-D2-GS | CAP-D2 (AT3G57060) | 3 | 4539 |
| CAP-D2-GS | SMC4 (AT5G48600) | 3 | 2543 |
| CAP-D2-GS | SMC2A (AT5G62410) | 3 | 2497 |
| CAP-D2-GS | SMC2B (AT3G47460) | 3 | 1932 |
| CAP-D2-GS | CAP-G (AT5G37630) | 3 | 1421 |
| CAP-D2-GS | CAP-H (AT2G32590) | 3 | 920 |
| CAP-D3-GS | CAP-D3 (AT4G15890) | 3 | 6668 |
| CAP-D3-GS | SMC4 (AT5G48600) | 3 | 1572 |
| CAP-D3-GS | SMC2A (AT5G62410) | 3 | 819 |
| CAP-D3-GS | CAP-G2 (AT1G64960) | 3 | 218 |
| CAP-D3-GS | CAP-H2 (AT3G16730) | 3 | 206 |
| CAP-D3-GS | SMC2B (AT3G47460) | 2 | 503 |

Among the proteins which co-purified with CAP-D2 (Table S1), other proteins such as the cohesin complex subunit SMC3 were identified. Additionally, the chromatin remodeling factors CHR17 and CHR19; CUL1, a subunit of the SCF ubiquitin ligase complex; HDC1, a histone deacetylase and ELO3, a histone acetyltransferase from the elongator complex were found. Among the proteins co-purifying with CAP-D3 (Table S2) were two nucleosome assembly proteins (NAP); CSN1, a subunit of the COP9 signalosome (CSN); the helicase BRAHMA; ELO3, from the elongator complex, and NERD, involved in DNA methylation.

The results indicate that both *A. thaliana Cap-D* genes are highly conserved, and that the corresponding proteins may act in combination with other condensin complex components, as well as with cohesin and SMC5/6 subunits.

### Condensin I subunits are localized within nuclei and cytoplasm

Previously, *A. thaliana* protoplasts have been used to examine the localization of the condensin subunits CAP-H and CAP-H2 (Fujimoto *et al.*, 2005). Therefore, we expressed transiently the coding region of *CAP-D2* fused to *EYFP* (35S∷CAP-D2_EYFPc) in *A. thaliana* mesophyll protoplasts. To visualize EYFP, the protoplasts were immunolabeled with anti-GFP antibodies. We identified CAP-D2 in the cytoplasm and the DAPI-counterstained nucleus (Fig. 4a). In the cytoplasm GFP-negative, but DAPI-positive round chloroplasts were also visible. The free EYFP of the positive control also localized in the cytoplasm and nucleus. Western blot analysis of CAP-D2_EYFPc transformed protoplasts confirmed that the CAP-D2_EYFP protein was intact, and that the visible localization corresponds to the fusion protein (187 kDa), and not to free EYFP (27 kDa) (Fig. 4b). The condensin I subunits, CAP-H and CAP-G fused to EYFP localized also in the cytoplasm and the nucleus (Figure S4). Similarly, transient transformation of *N. benthamiana* leaves revealed the localization of CAP-D2, CAP-H and CAP-G EYFP-fusion proteins in cytoplasm and nuclei too (Figure S5).

**Figure 4.**
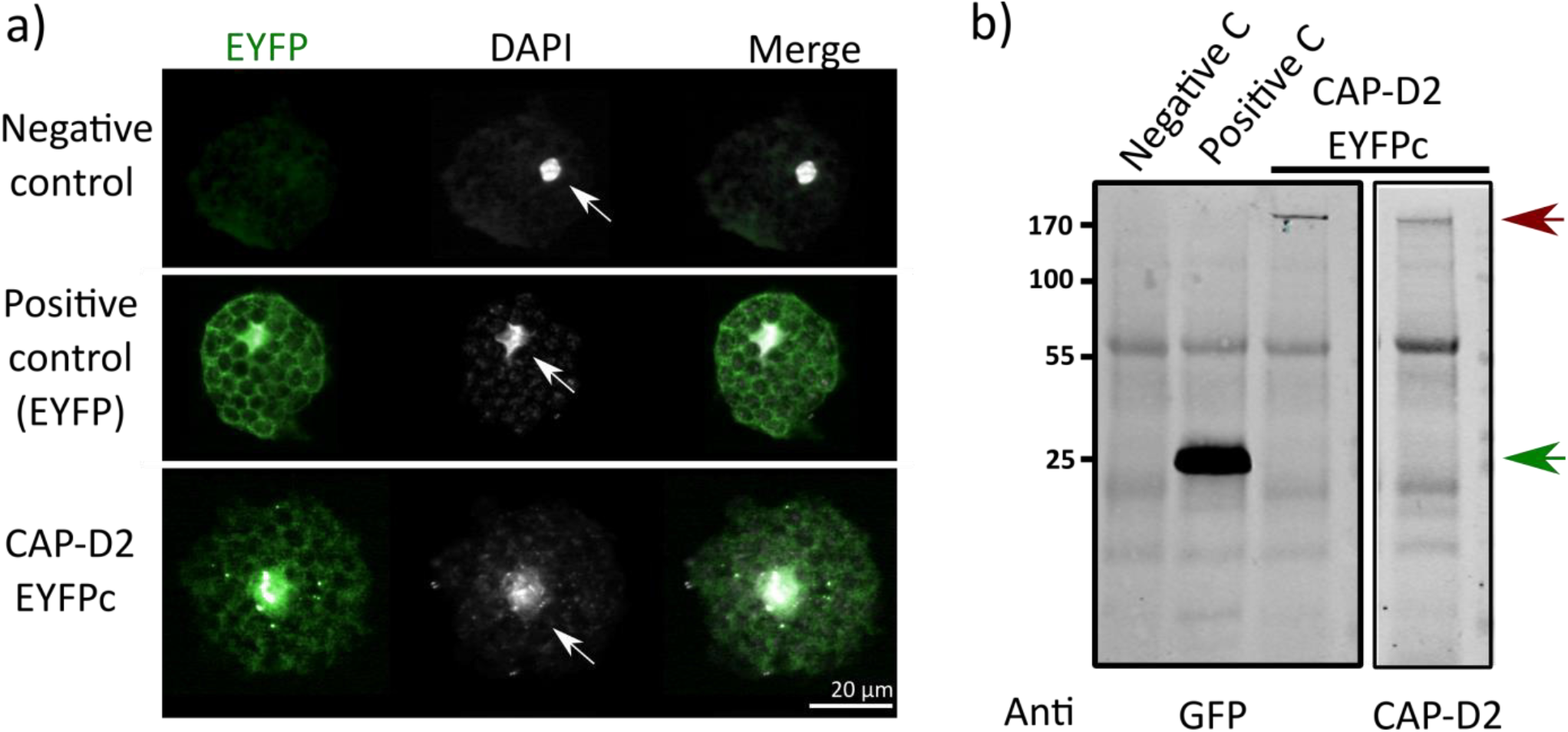
Localization of CAP-D2 in protoplasts. **a)** CAP-D2 is present in the cytoplasm and nuclei of leaf protoplasts. Untransformed protoplasts (negative control) and transformed with *CAP-D2* fused C-terminally to *EYFP*, and free *EYFP* (positive control) are presented. DAPI staining indicates the nuclei (arrows) and small signals in the cytoplasm corresponding to chloroplast DNA. **b)** Western blot analysis on total protein extracts from protoplasts untransformed (negative C), transformed with free *EYFP* (positive C) or with CAP-D2-EYFPc. The detection was performed with anti-GFP antibodies or anti-CAP-D2 serum. The intense band of 27 kDa (green arrow) corresponds to free EYFP. The bands of 187 kDa (red arrow) correspond to the CAP-D2_EYFP fusion protein.

Anti-CAP-D2 Western blot antibodies were generated against a recombinant protein containing the last 501 amino acids of CAP-D2 (Figure S6). The CAP-D2 antiserum can detect amounts as low as 1 ng of the recombinant protein (Figure S7). The CAP-D2 antiserum detects the CAP-D2 fusion protein from protoplasts (Fig. 4b), but not the CAP-D2 protein from wild-type leaves (data not shown). This may be due to a lower amount of the target protein in leaves compared to that in protoplasts. In protoplast overexpression of CAP-D2 occurred since the reporter construct is under control of the 35S promoter.

In order to localize CAP-D2 and CAP-D3 proteins *in planta*, *A. thaliana* wild-type plants were transformed with constructs containing the coding region of either gene fused at its C-terminus to enhanced yellow fluorescence protein (EYFP) under the control of the 35S promoter (*35S∷CAP-D2_EYFPc* and *35S∷CAP-D3_EYFPc*). In both cases, the detection of the proteins *in vivo* or by immunolocalization with anti-GFP antibodies (also detecting EYFP) was not possible. The same negative result was obtained by reporter constructs with EYPF fused at the N-terminus (*35S∷CAP-D2_EYFPn* and *35S∷CAP-D3_EYFPn*).

### CAP-D3 organizes chromatin during interphase

The involvement of *A. thaliana* CAP-D3 in compacting chromosome territories (CT) and keeping centromeres apart at interphase has been previously described by Schubert *et al.* (2013). In *Drosophila*, CAP-D3 is also involved in the formation of compact chromosome territories (Hartl *et al., 2008)*. To further study the involvement of CAP-D3 in chromatin organization we used two *cap-d3* mutants described previously, *Cap-D3* SAIL_826_B06 and *Cap-D3* SALK_094776 (Schubert *et al.*, 2013) (Fig. 5a,b). To confirm the centromeric clustering and CT dispersion phenotypes in both mutants, a FISH experiment on flow-sorted 4C nuclei was performed with probes specific for the centromere repeat pAL and the chromosome 1 arm territory bottom (CT1B) (Fig. 5c, d). In addition to the number of centromeric pAL signals per nucleus, the areas of the CT1B signals and the nucleus were measured. The median area size of the CT1B signals was 3.9, 4.7 and 4.7 μm^2^ for *cap-d3 SAIL*, *cap-d3 SALK* and wild-type, respectively (Fig. 5e). No significant differences were found. Thus, we could not confirm the CT dispersion phenotype of the *cap-d3* mutants described in Schubert *et al.* (2013). In addition, no significant differences were found in the nuclear area size between the *cap-d3* mutants and wild-type plants (Fig. 5e).

**Figure 5.**
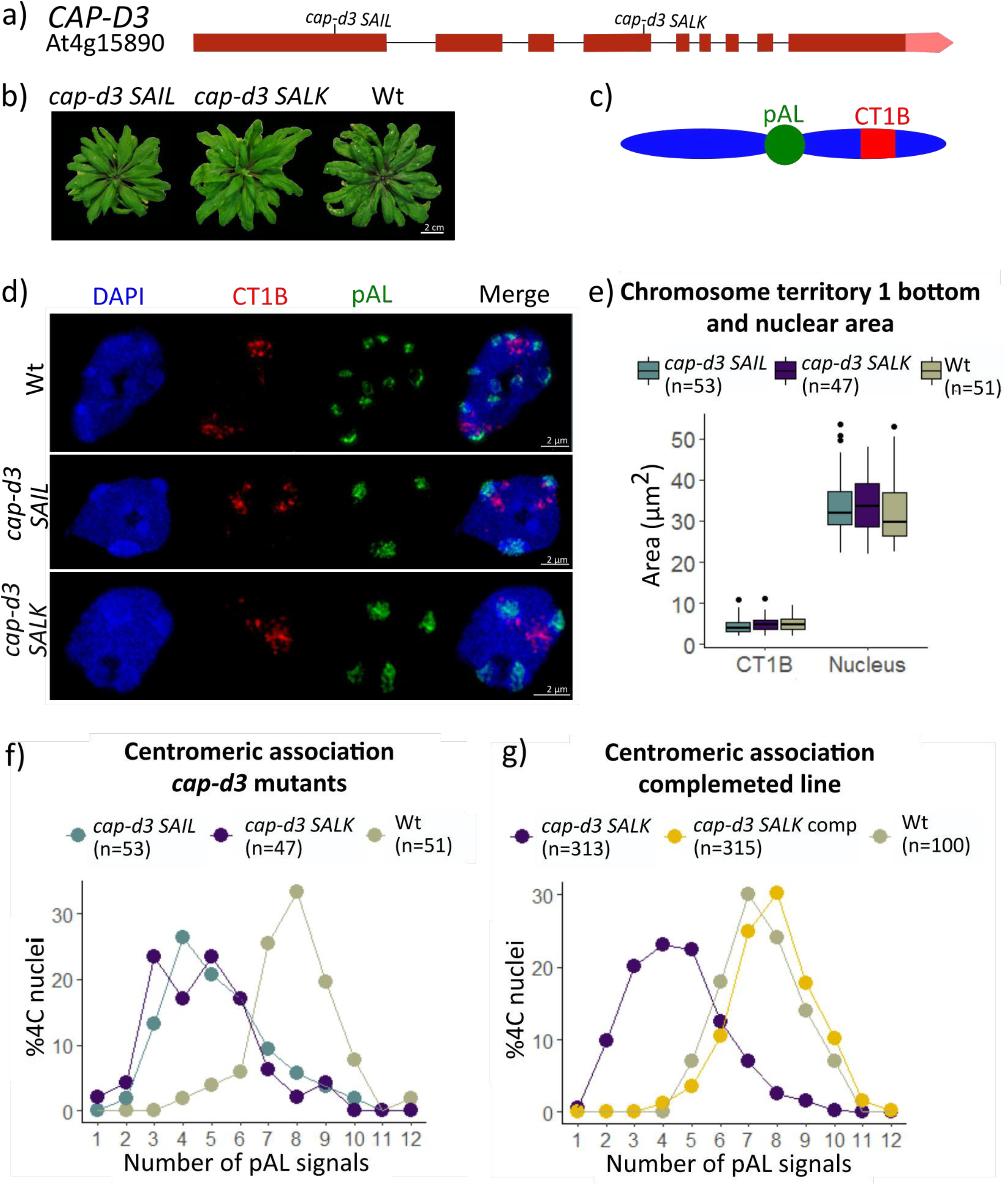
*cap-d3* mutations impair centromere distribution, but not CT compaction. **a)** Gene structure model of CAP-D3. Red boxes represent exons, lines the introns and the lighter red box the 3’UTR. The T-DNA insertion sites of the *cap-d3 SAIL* and *cap-d3 SALK* lines are indicated. **b)** Rosette leave stage phenotypes of homozygous *cap-d3 SAIL*, *cap-d3 SALK* and wild-type (Wt) plants. **c)** Schematic representation of chromosome 1 and the localization of the chromosome territory 1 bottom part (CT1B), and the centromeric pAL probe (labelling all ten centromeres present in the nuclei). **d)** SIM of a 4C nucleus labelled by FISH with the CT1B and pAL probes. **e)** Box plot diagram of the CT1B and the nucleus area sizes of *cap-d3 SAIL*, *cap-d3 SALK* and Wt nuclei. The boxes indicate upper and lower quartiles and the black bar the median. **f)** and **g)** pAL signal frequencies in 4C nuclei of *cap-d3 SAIL*, *cap-d3 SALK*, *cap-d3 SALK* complemented and Wt. n = total number of nuclei analyzed from two different plants in **e)** and **f)**, and from three different plants in **g)**.

On the other hand, we could confirm the centromere-association phenotype. In both *cap-d3* mutants the nuclei showed a lower number of centromeric pAL signal clusters than wild-type (Fig. 5d). Around 80% of the *cap-d3* mutant nuclei showed less than six pAL signals, while in wild-type only 12% of nuclei had less than six pAL signals (Fig. 5f). To verify that the mutation in the *CAP-D3* gene is indeed responsible for the centromeric clustering, a complementation experiment was carried out. *cap-d3 SALK* mutant plants were transformed with *CAP-D3_EYFPc* constructs, containing the coding region of *CAP-D3* fused to EYFP under the control of the 35S promoter. The centromeric association phenotype was evaluated in *cap-d3 SALK* complemented plants by FISH and compared with the *cap-d3 SALK* mutants and wild-type. Only 15% of the complemented nuclei showed less than six centromeric signal clusters, which is similar as the wild-type association levels (Fig. 5g). This confirms that CAP-D3 is responsible for the centromere association in the mutants.

Beside centromeres, in *A. thaliana*, the 45S and 5S rDNAs are heterochromatin-associated sequences. In nuclei of differentiated cells, 45S rDNA containing nucleolar organizing regions tend to associate, but the 5S rDNA loci are often separated (Berr and Schubert, 2007). To examine whether CAP-D3 affects in general the organization of heterochromatin, the distribution of the 45S and 5S rDNA loci was analyzed by FISH in both *cap-d3* mutants (Fig. 6a). The majority of 45S rDNA signals is shifted from three signals in wild-type to two signals in the mutants (Fig. 6b). No difference was observed with regard to 5S rDNA since over 70% of the nuclei showed between six and ten signals in the *cap-d3* mutant and wild-type plants (Fig. 6c). Thus, the *cap-d3* mutants present a higher association of the chromosomal 45S rDNA regions than wild-type, but the number of 5S rDNA signals remains unaffected.

**Figure 6.**
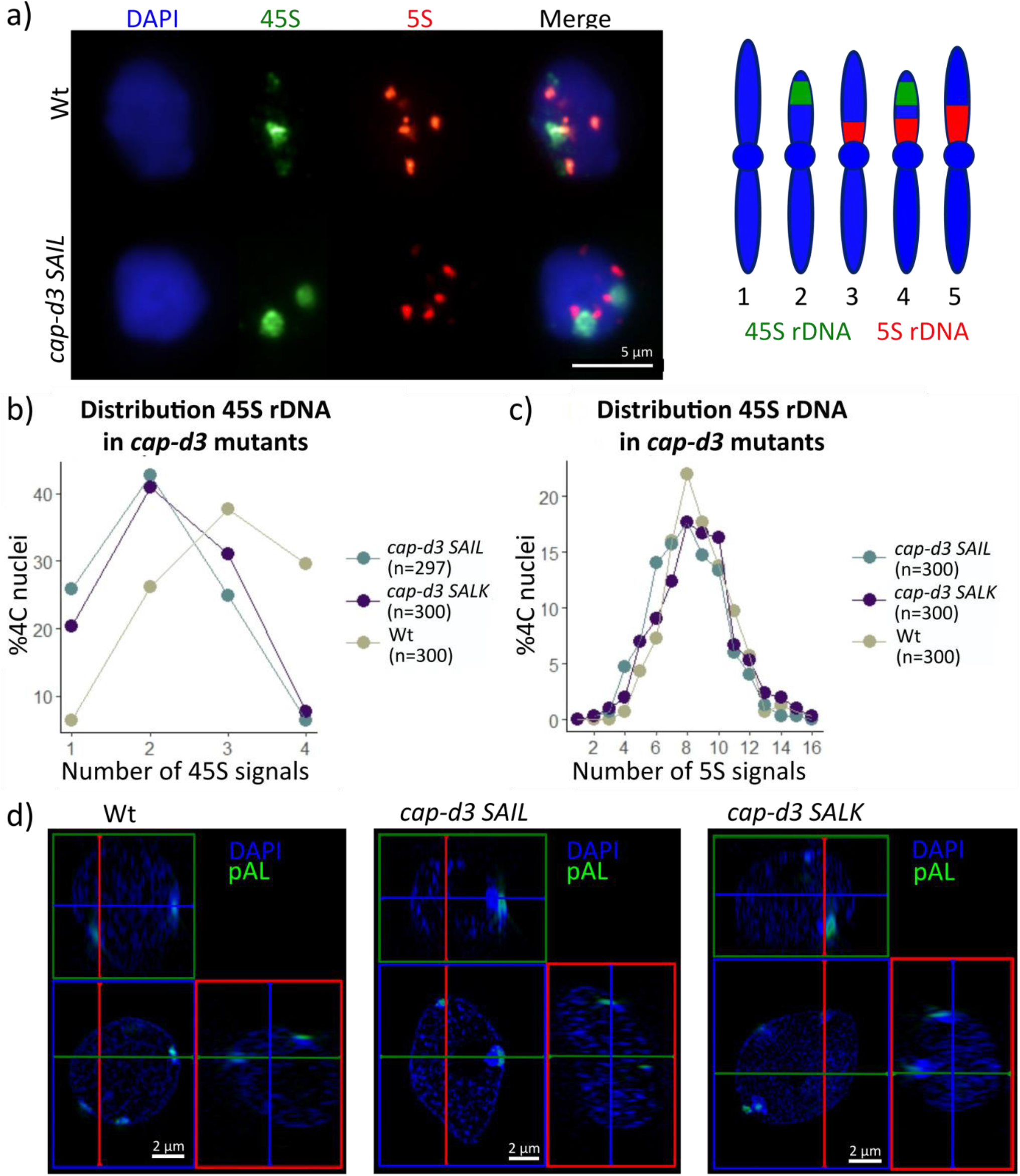
cap-d3 mutations induce of 45S rDNA association, but do not influence 5S rDNA and the spatial centromere arrangement in interphase nuclei. **a)** FISH signals of 45S rDNA and 5S rDNA on 4C nuclei of wild–type (Wt) and cap-d3 SAIL mutants. The ideogram (right) represents the *A. thaliana* chromosomes, showing the localization of 45S and 5S rDNA. **b)** and **c)** Frequency of 45S and 5S rDNA signals in 4C nuclei of Wt and the cap-d3 mutants. n = total number of nuclei analyzed from three different plants. **d)** SIM orthogonal view of FISH with the centromeric repeat (pAL) on structurally preserved acrylamide-embedded nuclei of Wt and cap-d3 mutants. Blue, green and red rectangles show x-y, x-z and y-z optical cross-sections, respectively.

*A. thaliana* centromeres are positioned at the nuclear periphery (Fransz *et al.*, 2002; Fang and Spector, 2005). To test whether the centromere position is influenced by the *cap-d3* mutations nuclei were embedded in acrylamide to preserve their 3D structure followed by FISH (3D-FISH) with the centromeric pAL repeats. For each genotype, *cap-d3 SAIL*, *cap-d3 SALK* and wild-type, 10 nuclei were analyzed. Optical sections (3D-SIM stacks) were acquired for each nucleus, and the centromere positions were analyzed in the ZEN software tool ‘ortho view’ (Fig. 6d). In all the cases, the centromeres were localized at the periphery of the nucleus, even when centromere clustering was present in the *cap-d3* mutants. Consequently, no deviation in peripheral centromere positioning in wild-type and the *cap-d3* mutants was observed.

### CAP-D3 does not effect the nuclear distribution of histone marks

DNA can be methylated at cytosine as 5-methyl-cytosine (5mC). The methylation of DNA is associated with heterochromatin formation and consequently, it has been found in the chromocenters of *A. thaliana* (Fransz *et al.*, 2002). Mouse embryonic stem cells depleted of condensin show a reduction of 5mC (Fazzio and Panning, 2010). In order to test whether such an effect can also be observed in plants, the distribution of methylated DNA in *cap-d3* mutants was compared to wild-type by immunodetection of 5mC-specific sites. In both *cap-d3* mutants and wild-type the 5mC signals were similarly chromocenter-localized (Fig. 7a). The *A. thaliana* centromeric repeats are highly methylated in a CpG context (Martinez-Zapater *et al.*, 1986). The use of methylation sensitive enzymes and Southern blot hybridization allowed a more precise determination of the relative DNA methylation level of the centromeric repeats. *HpaII* and its isoschizomer *MspI* cleave the same CCGG sequence, but *HpaII* is methylation sensitive while *MspI* is not. In wild-type, the centromeric repeats are highly methylated and are thus digestible by *MspI* (Fig. 7b). The ladder-like pattern corresponds to the monomer, dimer, trimer and higher orders of centromeric repeats. As expected, *HpaII* does not cut in wild-type DNA. In both *cap-d3* mutants, the hybridization pattern is similar to wild-type. Thus, the relative level of CCGG methylation is not altered in the *cap-d3* mutants (Fig. 7b).

**Figure 7.**
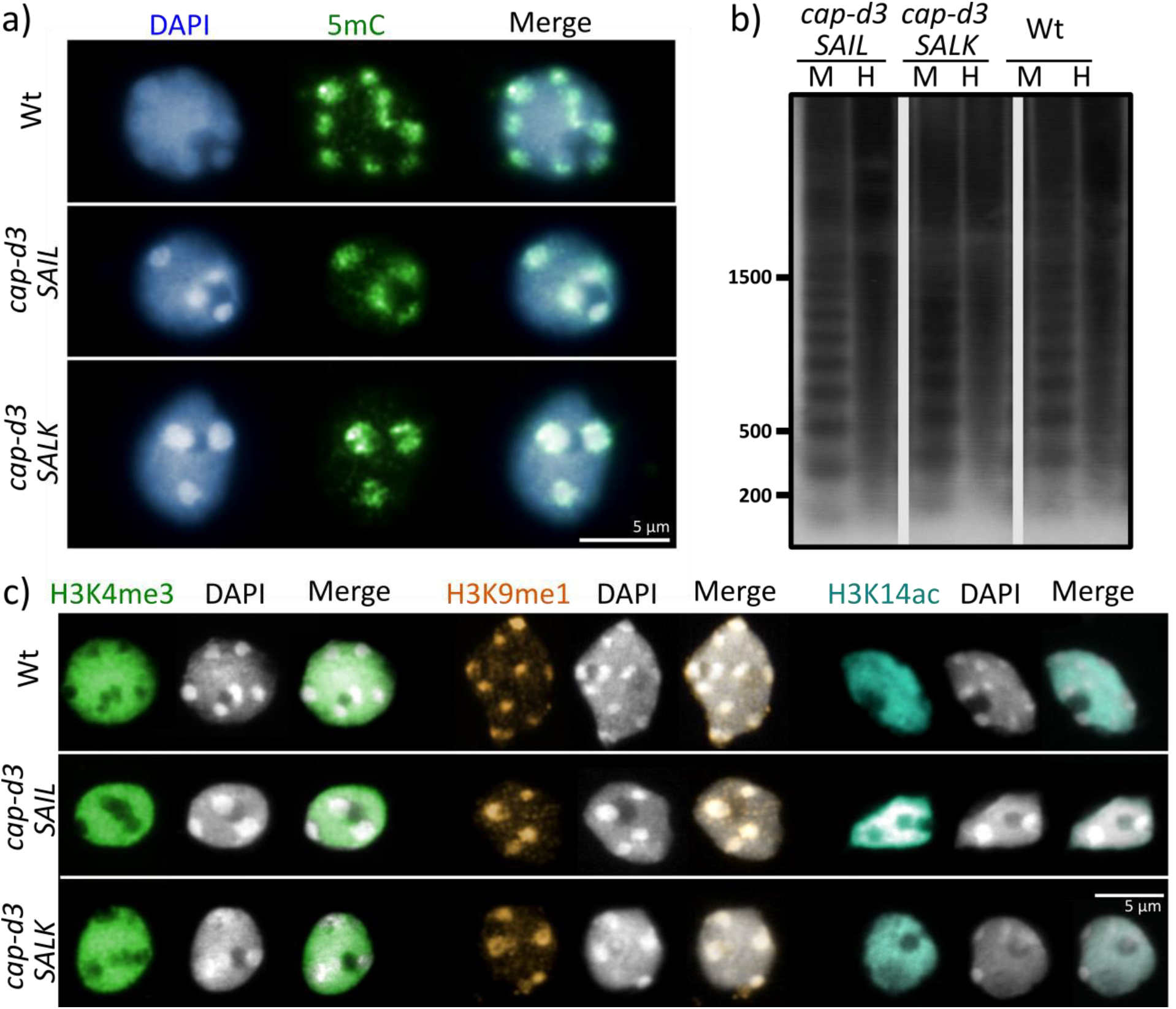
*cap-d3* mutations do not modify the epigenetic landscape in interphase nuclei. **a)** 5-methyl-cytosine immunolocalization on 4C nuclei of wild-type (Wt) and the *cap-d3* mutants. **b)** Southern blot analysis of the *cap-d3* mutants and Wt genomic DNA digested with *HpaII* (H) or *MspI* (M) and hybridized with the P^32^-labelled centromeric repeat pAL do not show different digestion patterns. **c)** Immunolocalization of histone H3K4me3, H3K9me1 and H3K14ac on 4C nuclei of Wt and the *cap-d3* mutants.

CAP-D3 prevents the clustering of heterochromatin, but the CAP-D3 protein itself localizes in euchromatic regions during interphase. Both types of chromatin are characterized by specific posttranslational histone modifications marks (Fuchs *et al.*, 2006). To evaluate a possible functional association between histone modifications and CAP-D3 functions, the global distribution patterns of different histone marks were compared between the *cap-d3* mutants and wild-type. Specific marks for heterochromatin (histone H3K9me1, H3K9me2) and euchromatin (histone H3K4me3, H3K27me3) were tested by indirect immunostaining. In addition, the H3 acetylation marks H3K9ac, H3K14ac, and H3K18ac as well as H3K9+14+18+23+27ac were evaluated. Histone acetylation relaxes chromatin allowing different protein complexes to access DNA (Wang *et al.*, 2014). Thus, histone acetylation is associated with transcription, and hypoacetylation with transcriptional repression. In flow-sorted 4C wild-type nuclei, H3K4me3 localizes in euchromatin and it is absent from chromocenters and the nucleolus. In *cap-d3* mutants the localization is identical. H3K9me1 is a heterochromatin-specific histone modification that localizes in the chromocenters in both *cap-d3* mutants and wild-type. Finally, the acetylation mark H3K14ac localizes mainly in euchromatin (transcriptionally active chromatin) of wild-type nuclei, but also in the mutants (Fig. 7c). The other histone modifications tested (H3K27me3, H3K9me2, H3K9ac, H3K18ac and H3K14+18+23+27ac) followed also the same pattern in wild-type and the *cap-d3* mutants (Figure S8). Thus, we did not detect obvious differences in the (sub-)nuclear distribution patterns of the different histone marks between wild-type and the *cap-d3* mutants.

### CAP-D3 moderately affects transcription

To assess if the increased clustering of the centromeric interphase chromatin in the *cap-d3* mutants affects gene transcription, the transcriptome of both *cap-d3* mutants was compared to wild-type. RNA-sequencing was performed in 4 samples (pooled 4 weeks-old plantlets) for each genotype. After differential expression analysis, we could observe alterations between the *cap-d3* mutants and wild-type transcriptomes. The genes with at least 2-fold change transcription and a pAdj ≤ 0.05 were considered as differentially expressed genes (DEG) between two genotypes (Fig. 8a). The smallest difference was observed between *cap-d3 SAIL vs*. *cap-d3 SALK* (74 DEG), and the highest between *cap-d3 SAIL vs.* wild-type (398 DEG). *cap-d3 SALK vs.* wild-type was intermediate (97 DEG)(Fig. 8b). Both *cap-d3* mutants show centromere and 45S rDNA clustering, but *cap-d3 SAIL* plants showed additional growth defects that are absent in *cap-d3 SALK* plants. To separate the individual effect of each allele, in further analysis only the DEG shared by both mutants when compared to wild-type were considered. These 83 genes, common to the *cap-d3* mutation independently of the specific alleles, are subsequently referred to as “*cap-d3* DEG” (Fig. 8b and Table S3). These genes are distributed along all chromosome arms (Fig. 8c). According to their Gene Ontology (GO) enrichment, the *cap-d3* DEGs are mainly involved in transcription, particularly in biological processes affecting the response to water, stimuli and stress (Table 2). In agreement with their role in transcription, 13 out of the 83 *cap-d3* DEG are transcription factors (Table S3). We conclude that the influence of CAP-D3 directly on transcription is moderate. However, the DEG involvement in plant response to stress, and the high proportion of transcription factors indicate that CAP-D3 may influence transcription indirectly.

**Table 2.** Gene ontology (GO) categories enriched in the 83 *cap-d3* differentially expressed genes (DEGs). FDR (False Discovery Rate): p-value adjusted for multiple testing.

| Ontology | GO term | Description | FDR |
| --- | --- | --- | --- |
| Biological process | GO:0009414 | Response to water deprivation | 0.0001 |
| Biological process | GO:0009415 | Response to water | 0.0001 |
| Biological process | GO:0042221 | Response to chemical stimulus | 0.0021 |
| Biological process | GO:0009628 | Response to abiotic stimulus | 0.0033 |
| Biological process | GO:0050896 | Response to stimulus | 0.01 |
| Biological process | GO:0006950 | Response to stress | 0.041 |
| Molecular function | GO:0030528 | Transcription regulator activity | 0.024 |
| Molecular function | GO:0003700 | Transcription factor activity | 0.047 |

**Figure 8.**
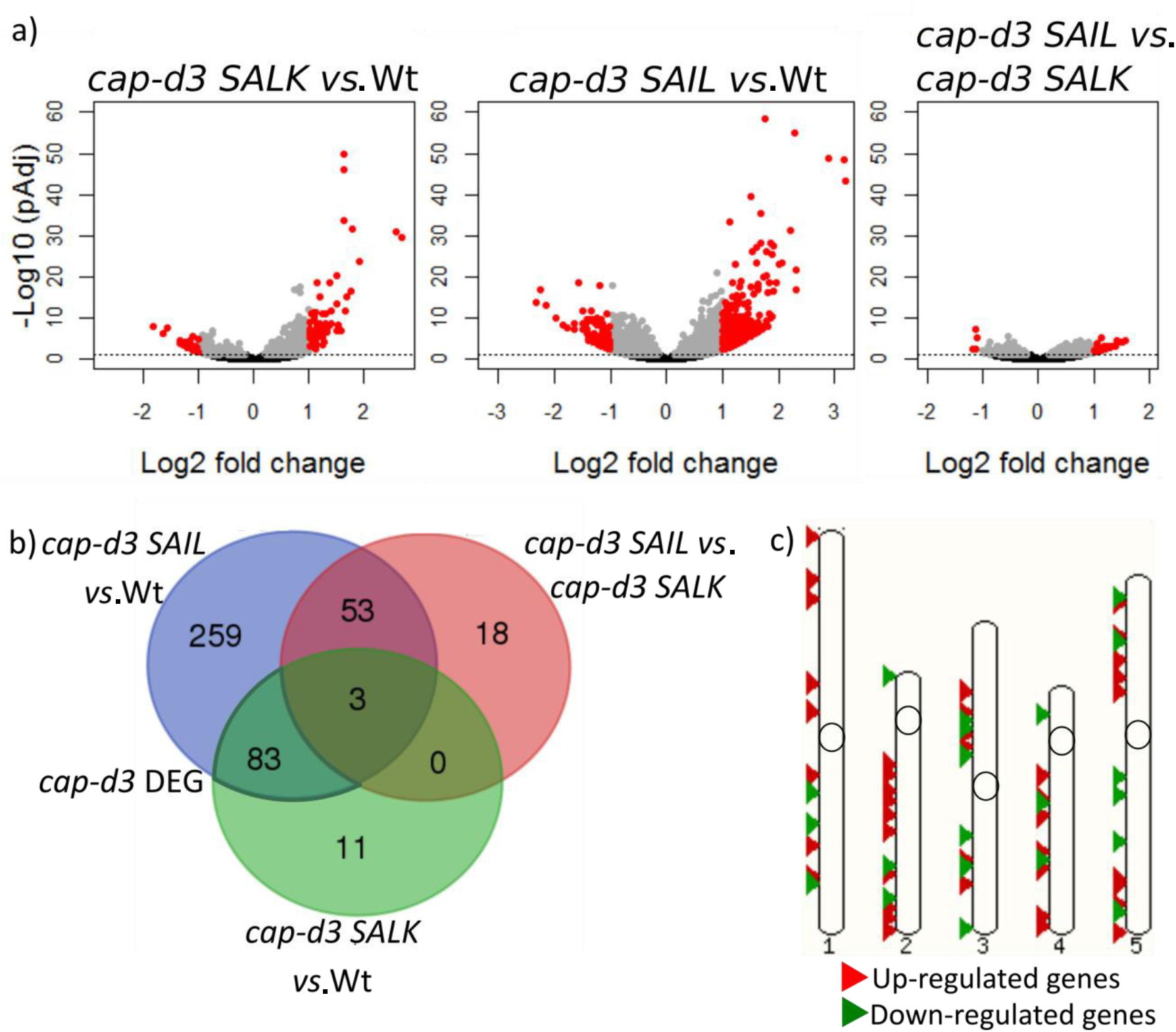
Transcriptome analysis of *cap-d3* mutants and wild-type plantlets. **a)** Volcano plots showing transcriptome comparisons between *cap-d3 SALK, cap-d3 SAIL* and Wt. The horizontal dotted line corresponds to pAdj = 0.05. Genes below are depicted in black and above in grey. The red genes are differentially expressed (DEG) at a threshold of 2 fold change (i.e., up-regulated: ≥ 1 Log2 FC, or down-regulated: ≤ −1 Log2 FC) and with a pAdj ≤ 0.05. pAdj is the p-value corrected for multiple testing with the Benjamini-Hochberg adjustment. **b)** Venn diagram showing the DEG across the three comparisons. Each circle comprises all the DEG genes of one comparison and the intersections between circles are the common DEG. For example: the blue circle represents the *cap-d3 SAIL vs.* Wt DEG, which are 398, of those: 83 are the same as in *cap-d3 SALK vs. Wt*, 53 are the same as in *cap-d3 SAIL vs. cap-d3 SALK*, 3 are differentially expressed in all comparison and 259 are only present in *cap-d3 SAIL vs.* Wt. **c)** The ideogram of *A. thaliana* chromosomes showing the position of the 83 *cap-d3* DEG along the chromosomes.

## DISCUSSION

### Arabidopsis CAP-D proteins are expressed in meristematic tissues in a cell cycle-dependent manner

*CAP-D2* and *CAP-D3* are highly expressed in meristems and cell cycle active tissues (flower buds, roots), but weaker in non-cycling tissues (mature leaves). Similarly, the condensin subunit genes *CAP-H* and *SMC2* are highly expressed in dividing tissues (Fujimoto *et al.*, 2005; Liu *et al.*, 2002; Siddiqui *et al.*, 2003). Sequences of 391 bp or 474 bp upstream of the start of *CAP-D2* or *CAP-D3*, respectively, are sufficient to act as promoters. Longer fragments (>1000 bp) do not improve the expression of the reporter gene. Interestingly, the *CAP-D2* and *CAP-D3* promoters regions contain two previously predicted E2F binding sites (Schubert *et al.*, 2013). E2F is a transcriptional activator of genes important for cell cycle progression. Together with retinoblastoma-related protein (RBR) and a dimerization partner, they control the transition from G1 to S phase. E2F sites are also present in the *A. thaliana* SMC2 promoter (Siddiqui *et al.*, 2003). In mouse, CNAP1 (CAP-D2) is also a target of E2F (Verlinden *et al.*, 2005). Considering the expression patterns, the promoter features and the comparison with other organism, it is plausible that the transcription of *A. thaliana CAP-D2* and *CAP-D3* is cell cycle-regulated.

Introns, when affecting the expression of a gene, often enhance its expression by increasing the transcript amount or by inducing the expression in specific tissues (Rose *et al.*, 2008; Parra *et al.*, 2011; Heckmann *et al.*, 2011). Nonetheless, the second intron of *CAP-D2* could have intragenic regulatory sequences repressing the expression in non-dividing tissues. This is supported herein by the loss of GUS reporter expression in leaves of the Pro6 and Pro9 transgenic plants compared with Pro5, Pro7 and Pro8 plants, which do not carry the second intron. Moreover, our quantitative RT-PCR results showed low transcription of *CAP-D2* in leaves. The second intron of the *AGAMOUS* gene is also responsible to inhibit expression in vegetative tissues, and drives its correct expression in flowers (Sieburth and Meyerowitz, 1997).

### The subunit composition of Arabidopsis condensin I and II is similar as in other eukaryotes

Protein immunoprecipitation (IP) from flower bud extracts confirmed already the presence of the subunits for condensin I and condensin II in *A. thaliana* (Smith *et al.*, 2014). Nonetheless, these IPs were performed with anti-SMC4, which would target both condensin complexes, and therefore could not determine the exact composition of condensin I and II. Our data based on affinity purification combined with mass spectrometry support that in *A. thaliana* both condensin complexes are present, and that their subunit composition is identical to those of other organisms (Hirano, 2012a). Notably, *A.thaliana* is the only species in which two SMC2 homologs have been predicted and described (Cobbe and Heck, 2004; Siddiqui *et al.*, 2003). Both, SMC2A and SMC2B can be active, but *SMC2A* accounts for most of the *SMC2* transcript pool (Siddiqui *et al.*, 2003). Both SMC2A and SMC2B interact with the other condensin subunits in vegetative and somatic tissues (Smith *et al.*, 2014; this study).

In human cells and *Drosophila*, CAP-D3 interacts with RBR and promotes the correct chromosomal localization of condensin II (Longworth *et al.*, 2008). In *A. thaliana*, this interaction is likely not conserved, since we could not detect RBR among the proteins that co-purified with CAP-D3. In human, Cdc20, a component of the anaphase-promoting complex E3 ubiquitin ligase, interacts and regulates CAP-H2 (Kagami *et al.*, 2017). In *Drosophila*, CAP-H2 also interacts and is regulated by the Skp cullin-F-box SCF^Slimb^ (Buster *et al.*, 2013), an E3 ubiquitin ligase regulated by CSN (Hotton and Callis, 2008). In our affinity purifications, we also detected components of the ubiquitin-26S proteasome pathway. CULLIN 1 co-purified with CAP-D2 and CSN1 with CAP-D3 in all replicates. CSN3 and CSN4 also co-purified with CAP-D3 in the three triplicates but also in 3 out of 115 of the non-specific proteins affinity purifications (data not shown). CULLIN1 and CULLIN3 were present in two of the CAP-D3 triplicates. These data suggest that in *A. thaliana*, ubiquitination could be involved in the regulation of the condensins.

A screen for functional partners of condensin in yeast identified, among others, two chromatin remodeling proteins and a histone deacetylase, as collaborators of condensin for chromosome condensation (Robellet *et al.*, 2014). In line with that, we identified chromatin remodeling enzymes (CHR17, CRH19 and BRAHMA), histone chaperones (NAP1;1 and NAP1;2), a histone deacetylase (HDC1) and a histone acetyltransferase (ELO3) in the affinity purification experiments with CAP-D2 and CAP-D3. All of them are chromatin modifiers important for plant development (Perrella *et al.*, 2013; Skylar *et al.*, 2013; Gentry and Hennig, 2014).

### Condensin I is located within the cytoplasm and nuclei during interphase

During interphase, in vertebrates the most commonly described localization of condensin I is exclusively in the cytoplasm (Hirota *et al.*, 2004; Ono *et al.*, 2004; Gerlich *et al.*, 2006; Hirano, 2012a). However, some reports regarding *Drosophila*, chicken and human cell cultures described the localization of condensin I additionally within the nucleus (Schmiesing *et al.*, 2000; Savvidou *et al.*, 2005; Zhang *et al.*, 2016). In *A. thaliana* protoplasts and *N. benthamiana* epidermal leaves, we observed CAP-D2, CAP-G and CAP-H EYFP-fusion proteins in the cytoplasm as well as the nucleus. The cytoplasmic localization of CAP-H was already described (Fujimoto *et al.*, 2005), but not yet its nuclear localization.

In stable *A. thaliana* transformants carrying CAP-D2 or CAP-D3 tagged at its N- or C-terminus to EYFP and in stable *cap-d3* mutants carrying CAP-D3-EYFP, the fusion proteins could not be visualized, neither directly nor indirectly. The constructs are functionally active, since they work in *A. thaliana* and in *N. benthamiana* after transient transformations. In addition, the CAP-D3-EYFP construct was able to complement the centromeric phenotype of the *cap-d3 SALK* mutants. Similar problems have been described for GFP-PATRONUS1 *A. thaliana* transformants (Zamariola et al., 2014). These authors suggested that the reason behind could be the low expression or stability of the PATRONUS protein due to the presence of an APC/C degradation box. However, in CAP-D2 no APC/C degradation box exists. The detection of CAP-D2 in leaves from *A. thaliana* wild-type plants by Western blot was also not possible. This may be due to a low protein level in wild-type leaves since the transcript level in leaves is very low. By Western blot the CAP-D2 protein was detectable in protoplasts only when constitutively overexpressed. Similarly, in *Drosophila* the detection of condensin from extracts of non-dividing tissues was also not possible (Cobbe *et al.*, 2006).

### CAP-D3 may influence interphase chromatin arrangement and transcription, but not histone modifications

In *Drosophila*, CAP-D3 and CAP-H2 are needed to form compact chromosomes (Hartl *et al.*, 2008; Bauer *et al.*, 2012). Condensins via maintaining chromatin condensation may also maintain nuclear shape and size, as indicated after SMC2, CAP-H2 and CAP-D3 depletion in human cells (George *et al.*, 2014). In embryonic stem cells of mice, the depletion of SMC2 causes chromatin decondensation as well as the increase of the nuclear volume (Fazzio and Panning, 2010). On the other hand, in *C. elegans*, the depletion of SMC4, CAP-G2 or HCP-6 (CAP-D3) does not change the chromosome territory volumes (Lau *et al.*, 2014).

In *A. thaliana*, previous studies based on FISH suggested an influence of CAP-D3 on the formation of the top arm 1 interphase chromosome territories and sister chromatid cohesion (Schubert *et al.*, 2013). Using FISH probes against a smaller part of chromosome 1 bottom arm, we could not detect an increase of the hybridization signal area in the *cap-d3* mutants compared to wild-type plants. The differences could be explained by labeling only one fourth of the chromosome arm by FISH, while in the previous study the whole chromosome arm (without pericentromeric heterochromatin) was visualized. The different methods used to quantify the dispersion of the interphase chromatin could be another reason. The degree of chromatin condensation within nuclei may depend on the type of tissue (Tessadori et al., 2007; van Zanten *et al.*, 2011). Light (Bourbousse *et al.*, 2015), drought, temperature, and salinity stress, as well as toxic components, energy-rich radiation and chemically induced DNA damage may also induce dynamic structural changes in plant chromatin (reviewed in Probst & Mittelsten-Scheid, 2015). Even compressive stress has the potential to reorganize chromatin (Damodaran *et al.*, 2018; Xia *et al.*, 2018). Thus, these factors have also the potential to influence chromatin condensation in the CAP-D mutants.

Although we could not confirm the euchromatin dispersion in the *cap-d3* mutants, it cannot be excluded that CAP-D3 is involved in the organization of chromosome territories as found in *Drosophila* (Bauer *et al.*, 2012; Hirano, 2012b). The mutants used in our analysis (*cap-d3 SAIL* and *cap-d3 SALK*) have knockdown alleles, meaning that there is still a truncated transcript that could produce a partially functional CAP-D3 protein (Schubert *et al.*, 2013).

In addition to its role in chromosome compaction, condensin II has been described to influence transcription (Longworth *et al.*, 2012; Dowen *et al.*, 2013; Yuen *et al.*, 2017). Although the *A. thaliana cap-d3* mutants showed only moderate transcriptional changes, CAP-D3 might still affect the expression of genes involved in transcription and response to stress. This conclusion arises from our observation on the *cap-d3* mutants which die sooner than wild-type plants after stress, such as pathogen infection. Interestingly, gross chromosome rearrangements altering the genome topology do not alter gene expression in *Drosophila* (Ghavi-Helm *et al.*, 2018). Even a budding yeast strain, after merging all 16 chromosomes into a single one, revealed a nearly identical transcriptome and similar phenome profiles as wild-type strains (Shao *et al.*, 2018). Thus, chromatin structure changes as induced in the *A. thaliana cap-d3* mutants seem to influence the global transcription only slightly.

Wang *et al.* (2017) showed that *A. thaliana* SMC4, but not CAP-D3, is important to maintain the repression of pericentromeric retrotransposons independent of DNA methylation. Accordingly, we observed no increased retrotransposon transcription in any of the *cap-d3* mutants. Moreover, in accordance with our observations for both *cap-d3* mutants, the protein-coding genes up-regulated in *smc4* mutants are mainly involved in flower development, reproductive processes and DNA repair, and are distributed all over the genome (Wang *et al.*, 2017). However, we observed in the *cap-d3* mutants a differential expression of genes involved in transcription and stress response. This difference could be due to the combined effects of both condensin complexes I and II in the *smc4* mutants, while in our *cap-d3* mutants only condensin II is compromised.

Posttranslational histone modifications may affect the structure and stiffness of interphase nuclei, and decondensed euchromatin correlates with less rigid nuclei (Chalut *et al.*, 2012; Krause *et al.*, 2013; Haase *et al.*, 2016). In human HeLa cells histone methylation, but not acetylation, contributes to the stiffness and structure of condensed mitotic chromosomes (Biggs *et al.*, 2019). Histone acetylation relaxes chromatin allowing different protein complexes to access DNA. Thus, acetylation is associated with transcription, and hypoacetylation with transcriptional repression (Wang *et al.*, 2014).

It seems that the unaltered degree and pattern of histone acetylation reflects an only moderate effect on transcription as we observed in the *capd-3* mutants.

### CAP-D proteins prevent heterochromatin clustering

CAP-H2 promotes the spatial separation of heterochromatic regions in *Drosophila* during interphase (Bauer *et al.*, 2012; Buster *et al.*, 2013). Correspondingly, in *A. thaliana*, depletion of CAP-D3 results in centromere clustering at interphase (Schubert *et al.*, 2013). We confirmed this interphase phenotype and found that CAP-D3 depletion also results in the clustering of the 45S rDNA loci but not of the 5S rDNA sites. A differential behavior of rDNA was also found in protoplasts of *A. thaliana*. 45S rDNA remains condensed while the 5S rDNA decondenses during protoplast formation (Tessadori *et al.*, 2007). 5S and 45S rDNA are transcribed by RNA polymerases III and I, respectively (Layat *et al.*, 2012). Therefore, the different clustering behaviors of both rDNAs in the *cap-d3* mutants could be due to their different structural and functional properties. Moreover, condensin of fission yeast, which is similar to condensin I, binds to RNA polymerase III transcribed genes (5S rDNA and tRNA), and mediates their localization near the centromeres (Iwasaki *et al.*, 2010).

The nuclear and chromocenter phenotype which we observed in the *cap-d3* mutants differs from previous reports (Moissiard *et al.*, 2012; Sakamoto and Takagi, 2013; Tatout *et al.*, 2014; Poulet *et al.*, 2017; Wang *et al.*, 2017). The chromocenters cluster and localize at the nuclear periphery, but do not decondense, the nuclear area does not change compared to that of wild-type, and the general degree of DNA and histone methylation is unaffected. Moreover, hypomethylation, *linc* and *morc* mutants do not show transcriptional silencing of centromeric and pericentromeric repeats, and of silenced genes (Moissiard *et al.*, 2012; Poulet *et al.*, 2017). In contrast, CAP-D3 has little effect on silencing, because no increased transcription of transposable elements was detected in *cap-d3* mutants (Wang *et al.*, 2017). MORC, CRWN and LINC proteins localize close to the chromocenters, MORC foci adjacent to the chromocenters (Moissiard *et al.*, 2012), CRWN1 and CRWN4 at the nuclear periphery (Sakamoto and Takagi, 2013), and the LINC complex in the nuclear envelope (Tatout *et al.*, 2014). Conversely, CAP-D3 influences the arrangement of the chromocenters but localizes exclusively in euchromatin during interphase (Schubert *et al.*, 2013). Therefore, CAP-D3 has mainly a structural role during interphase and affects the clustering of chromocenters without localizing close to them.

Statistical analysis detected a more regular, than a completely random spatial centromere and chromocenter distributions in animal and plant nuclei. This suggests that repulsive constraints or spatial inhomogeneities influence the 3D organization of heterochromatin (Andrey *et al.*, 2010). Computer simulation modeling of *A. thaliana* chromosomes as polymers predicts that the position of the chromocenters in the nucleus is due to non-specific interactions (de Nooijer *et al.*, 2009). The simulated chromosomes exhibit chromocenter clustering except for the so-called Rosette chromosomes, in which the euchromatin loops emanate from the chromocenter and thus prevent chromocenter clustering (Fransz *et al.*, 2002). Indeed, depletion-attraction forces predict that big particles in an environment crowded with small particles will tend to cluster together (Marenduzzo *et al.*, 2006). This situation can be applied to the nucleus where the chromocenters act as big particles and euchromatin as small particles. If association is not prevented, the chromocenters will cluster.

In *cap-d3* mutants we observed chromocenter clustering but barely chromosome territory dispersion. During mitosis, CAP-D3 is needed to confer the rigidity of the chromosome arms (Green *et al., 2012)* and human condensin controls the elasticity of mitotic chromosomes (Sun *et al.*, 2018).

We suppose, that during interphase, CAP-D3 localizes in euchromatin, possibly along the euchromatic loops, mediating the rigidity which is needed to keep the chromocenters away from each other. In case of lacking or functionally impaired CAP-D3, the loops may be not stiff enough to prevent the chromocenter clustering while the chromosome territories may mainly keep their structures (Fig. 9). The finding that condensed chromatin resist to mechanical forces, whereas decondensed chromatin is more soft (Maeshima *et al.*, 2018) supports the idea that the stiffness of chromatin is an important feature to organize cell nuclei. Our observation that the degree of methylation and acetylation is not altered in the *cap-d3* mutants suggests that these post-translational histone modifications are not required for the rigidity of interphase chromosome territory structures.

**Figure 9.**
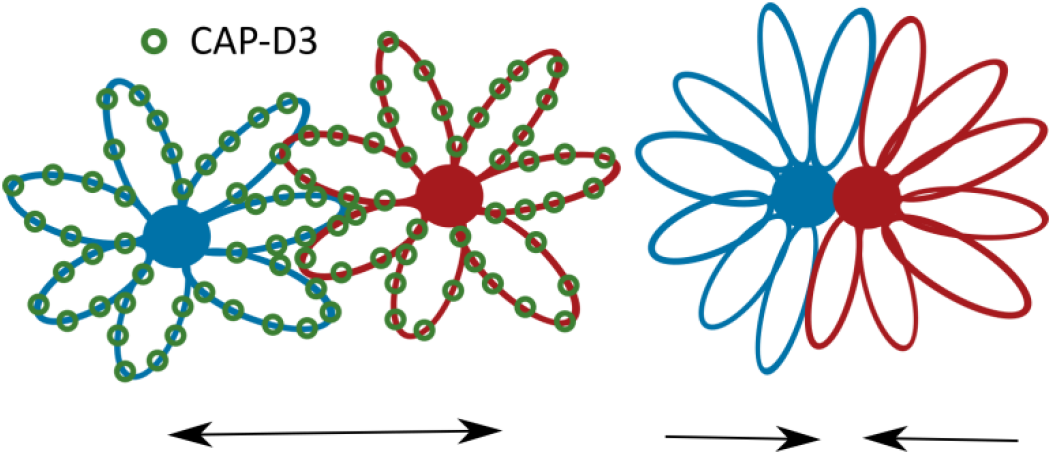
Model explaining the function of *Arabidopsis* CAP-D3 in interphase nuclei. Two chromosomes are represented in blue and in red with euchromatin emanating loops from their pericentromeric chromocenters (rosette chromosome model; Fransz et al., 2002; de Nooijer et al., 2009). CAP-D3 (green circles) localizes along euchromatin creating the chromatin loops rigid to keep the chromocenters separated (left). In absence of CAP-D3 the chromatin loops are not stiff enough to counterbalance the depletion-attraction forces (Marenduzzo et al., 2006). Consequently, the chromocenters cluster (right).

## EXPERIMENTAL PROCEDURES

### Plant material and stable transformation

All *Arabidopsis thaliana* (L.) Heynh lines and control plants are in Columbia-0 (Col-0) background. The T-DNA *cap-d3* (SAIL_826_B06, SALK_094776) insertion lines were previously described and selected in our laboratory (Schubert *et al.*, 2013). Seeds were sown in soil and germinated under short-day conditions (16 h dark/8 h light, 18-20°C) and then transferred to long-day conditions (16 h light/ 8 h dark, 18-20°C) before bolting. The lines were genotyped by PCR using the primers listed in Table S4. The presence of the T-DNA was further confirmed by sequencing.

*A. thaliana* stable transformants were generated by the floral dip method (Clough and Bent, 1998). For selection of primary transformants, the seeds were sterilized and plated on ½ Murashige and Skoog (MS) basal medium (Sigma) supplemented with the adequate antibiotics when required and grown in a growth chamber under-long day conditions.

### Transcript quantification

For transcript quantification total RNA was extracted from leaves, roots, 7 days-old seedlings and flower buds with the RNeasy Plant Mini kit (Qiagen) following manufacturer’s instructions. All RNA samples were treated with TURBO DNAse (Thermo Fisher Scientific) and tested for DNA contamination by PCR. Reverse transcription was performed using 250 ng of total RNA and the RevertAid H Minus First Strand cDNA Synthesis kit (Thermo Fischer Scientific), with oligo(dT)18 primers, according to manufacturer’s instructions. The quality of the cDNA was checked with a PCR targeting EF1B mRNA (Elongation factor 1β).

Quantitative RT-PCRs for *CAP-D2* and *CAP-D3* transcripts were done in triplicates and from three independent biological samples using SYBR™ Green PCR Master Mix (Thermo Fischer Scientific) in a 7900HT Fast Real-Time PCR System (Applied Biosystems). For each reaction, 0.5 μl of cDNA template and 0.6 mM primers (Table S4) were used in 10 μl. *PPA2* and At4g26410 (Kudo *et al.*, 2016) were used as reference genes for data normalization and the data were analyzed with the Double Delta Ct method (Livak and Schmittgen, 2001).

### *CAP-D2* and *CAP-D3* promoter∷GUS reporter lines and β-glucuronidase activity assay

Different lengths of the promoter regions of both *CAP-D2* and *CAP-D3* were cloned between the *Sal*I and *Not*I restriction sites of the pEntr 1A plasmid (Invitrogen). The sequences were amplified from gDNA with the primer pairs D2-1156F/D2ProR for the Pro4 fragment, D2-1156F/D2Int1R for Pro5, D2-1156F/D2Int2R for Pro6, D2-392F/D2ProR for Pro7, D2-392F/D2Int1R for Pro8, D2-392F/D2Int2R, for Pro9, D3-1318F/D3ProR for Pro10 and D3-474F/D3ProR for Pro11 (Table S4). The fragments were subcloned upstream of the GUS reporter gene in the pGWB633 plasmid (Nakamura *et al.*, 2010) using the Gateway LR Clonase II enzyme mix (Invitrogen) following manufacturer instructions.

Constructs were transformed into *A. thaliana* and stable transformants were selected in ½ MS (Sigma) with 16 mg/L PPT (Duchefa). One month after sowing, the plantlets were stained for GUS activity according to Jefferson *et al.* (1987) with small modifications. Plantlets were collected in 15 ml tubes containing 1% X-Glu (5-Bromo-4-Chloro-3-indolyl-β-D-Glucopyranoside; Duchefa) in 0.1 M phosphate buffer (pH 7.0). To facilitate the penetration of the solution in the material, the tubes with the plant material and the staining solution were exposed to vacuum for 5 min and incubated overnight at 37°C. Next day, the staining solution was replaced by 70 % ethanol and incubated 20 min at 60°C. This step was repeated until the chlorophyll was removed. The stained material was preserved in 70% ethanol at 4C° and analyzed under a stereo microscope.

### Condensin subunit EYFP-fusion constructs

The 3942 bp and 4245 bp long cDNA sequences of *CAP-D3* and *CAP-D2* respectively, were synthesized and cloned into pEntr 1A (Invitrogen) by a DNA-Cloning-Service (Hamburg, Germany). The 3153 bp and the 2013 bp long cDNA sequences of *CAP-H* and *CAP-G* were amplified from flower bud cDNA with the primer pairs CAPH_pentry_f/CAPH_pentry_r and CAPG_pEnt_f/CAPG_pEntr_r (Table S4), respectively, and cloned between the *Sal*I and *Not*l sites of the pEntr 1A plasmid (Invitrogen).

Once in the pEntr 1A plasmid, the coding sequences of the genes of interest were subcloned into pGWB641 and pGWB642 plasmids (Nakamura *et al.*, 2010) using Gateway cloning (Invitrogen). The generated expression cassettes contained the proteins of interest fused to EYFP C-terminally for the pGWB641 constructs (CAP-D2_EYFPc, CAP-D3_EYFPc, CAP-G_EYFPc and CAP-H_EYFPc) or N-terminally for the pGWB642 constructs (CAP-D2_EYFPn and CAP-D3_EYFPn), and both were under the control of the cauliflower mosaic virus 35S promoter. As a control (Control_EYFPc), a plasmid containing only EYFP under the control of the 35S promoter was generated.

### Condensin I subunit localization in *A. thaliana* protoplasts and *N. benthamiana*

Isolation and transformation of *A. thaliana* leaf protoplasts were performed as described in Yoo *et al.*, (2007). To improve the visualization of the fusion proteins, the transformed protoplasts were fixed in 4% formaldehyde in 1×PBS, washed in 1×PBS and centrifuged at 400 rpm for 5 min (Shandon CytoSpin3, GMI inc) onto a microscopic slide. The slide was directly used for immunostaining against EYFP.

*Nicotiana benthamiana* leaf cells were transformed as described in Sparkes *et al.*, (2006). Agrobacteria carrying the constructs of interest were grown in YEB medium with suitable antibiotics to an OD_600_ of 1 and resuspended in infiltration medium (10 mM MES, 10 mM MgCl_2_, pH 5.6, 3.3 mM acetosyringone). *N. benthamiana* leaves of 2 to 3 weeks old plants were infiltrated with the Agrobacterium suspension using a syringe without needle and analyzed 2 to 4 days later.

### Antibody production

The sequence between 2743 and 4248 bp of *CAP-D2* (501 C-terminal amino acids) was amplified from *A. thaliana* flower bud cDNA with the D2CtSalI_F and D2CtNotlI_R primers (Table S4). The fragment was cloned between the *Sal*I and *Not*I restriction sites of the pET23a(+) plasmid (Novagen) resulting in a pEt23_CAP-D2_Ct construct which contains the cassette T7 promoter∷T7 tag-*CAP-D2*Ct-His tag. The construct was transformed into *E. coli* BL21 cells and the expression of the transgene induced with 1 mM IPTG (isopropyl-β-D-thiogalatopyranoside, Sigma-Aldrich). The recombinant protein was purified with agarose beads that bind specifically to the His-tag (Ni-NTA Agarose, Qiagen) following the purification hybrid method from the ProBond purification system (Thermo Fisher Scientific). The purified recombinant protein was used to produce anti-CAP-D2 polyclonal antibodies in rabbit (Udo Conrad, Phytoantibody group, IPK, Gatersleben, Germany). Two rabbits were immunized with the recombinant proteins and the anti-serum collected after four immunizations. The anti-CAP-D2 serum was used directly for Western blot.

### Total protein extraction and Western blot

Isolated protoplast or grinded plant leaf material were resuspended in 100-300 μl of protein extraction buffer (56 mM Na_3_CO_3_, 56 mM DTT, 2% SDS, 12% Sucrose, 2 mM EDTA, bromophenol blue), incubated 20 min at 65°C and centrifuged at high speed (13,000 rpm). Then, the supernatant containing the soluble total protein was used for Western blot analysis.

For Western blot, the extracted proteins were separated in 10% polyacrylamide gels **(Schägger and von Jagow, 1987**) and blotted onto Immobilion-Fl membranes (Millipore). The membranes were incubated in blocking solution (5% skim milk in TBST) for 30 min to reduce non-specific binding of the antibodies. Then, the membranes were incubated overnight at 4°C with the adequate primary antibodies in TBST buffer: mouse anti-His-tag (1:2000, Millipore, 05-949), rabbit anti-CAP-D2 serum (1:1000) or rabbit anti-GFP conjugated with Alexa488 (1:1000, Chromotek, PABG1). And then with secondary antibodies: anti-mouse IgG IRDye 680RD (1:10000, LI-COR, 926-32222) or anti-rabbit IgG IRDye800CW (1:5000, LI-COR, 925-32213). The membranes were imaged using a LI-COR Odyssey Imager (LI-COR). For T7-tag detection, the membranes were first incubated with anti-T7-tag conjugated with alkaline phosphatase (1:10000, Merck, #6999) for 1 h, and then with phosphate-activity buffer (100mM Tris, 100 mM NaCl, 1 mM MgCl_2_, 0.33 mg/ml nitro blue tetrazolium and 0.17 mg/ml 5-bromo-4-chloro-3-indolyl phosphate) in the dark until the signals were visible.

### Affinity purification and analysis of GS-tagged CAP-D2 and CAP-D3 from PSB-D cells

The cDNA sequences of *CAP-D3* and *CAP-D2* were synthesized and cloned into pCambia 2300 35S GS-Ct by the DNA-Cloning-Service (Hamburg, Germany) resulting in the constructs pCambia2300_CAP-D2_GS and pCambia2300_CAP-D3_GS.

The *A. thaliana* ecotype ‘Landsberg erecta’ cell suspension (PSB-D) was transformed as described (Van Leene *et al.*, 2011). CAP-D2-GS and CAP-D3-GS were affinity purified following the protocol described (Dürr *et al.*, (2014).

For mass spectrometry, the eluted proteins were separated in a 10 % polyacrylamide gel and digested with trypsin. Mass spectrometry and data analysis were performed according to Antosz *et al.*, (2017). Protein Scape 3.1.3 (Bruker Daltonics) in connection with Mascot 2.5.1 (Matrix Science) facilitated database searching of the NCBInr database.

Three independent affinity purifications were performed. A MASCOT score of minimum 100 and the presence in at least two of the purifications were considered as criteria for reliable protein identification. The experimental background (contaminating proteins that co-purified with the unfused GS-tag) and non-specific interactions (proteins that co-purified independently of the bait used) were subtracted. The list of non-specific *A. thaliana* proteins is based on 543 affinity purification experiments using 115 different baits (Van Leene *et al.*, 2014).

### Nuclei preparations

*A. thaliana* nuclei from differentiated leaf cells were isolated and flow-sorted according to their ploidy level as described (Weisshart *et al.*, 2016) in a BD INFLUX Cell Sorter (BD Bioscience). The nuclei were sorted based on their DNA content in 2C, 4C, 8C and 16C ploidy fractions. Twelve μl of 4C sorted nuclei and the same amount of sucrose buffer (10 mM Tris, 50 mM KCl, 2 mM MgCl-6H_2_O, 5% sucrose, pH 8.0) were placed on a slide. The slides were directly used or stored at −20°C.

*A. thaliana* nuclei were embedded in acrylamide to preserve their 3D structure following the procedure described by Kikuchi *et al.*, (2005) with modifications. Twelve μl of nuclei suspension were mixed on a slide with 6 μl of active 15% acrylamide embedding medium (15% acrylamide/bisacrylamide (29:1), 15 mM PIPES, 80 mM KCl, 20 mM NaCl, 2 mM EDTA, 0.5 mM EGTA, 0.5 mM spermidine, 0.2 mM spermine, 1 mM DTT, 0.32 M sorbitol, 2% APS and 2% Na_2_SO_3_). A coverslip was carefully placed on top of the acrylamide-nuclei mixture and let to polymerize 30 min at room temperature. The coverslip was then removed letting a thin pad of nuclei embedded in acrylamide on the slide that was directly used for FISH.

### Preparation of squashed *A. thaliana* roots

*A. thaliana* seedlings were fixed in 4% paraformaldehyde in phosphate-buffered saline buffer (1×PBS buffer). The seedlings were washed in 1×PBS buffer and digested for 30 min at 37°C in an enzyme mix (0.5% pectolyase (Sigma), 0.5% cytohelicase (Sigma), 0.35% cellulase (Calbiochem), 0.35% cellulase (Duchefa) in 1×PBS buffer. After removal of the enzyme solution and washing in 1×PBS, the root tips were transferred to a slide and squashed between coverslip and slide. The liquid nitrogen frozen coverslip was lifted and the slide directly used for immunostaining.

### Probe preparation and fluorescence *in situ* hybridization (FISH)

The probes were generated by: (i) PCR for the 180 bp centromeric repeat (pAL; Martinez-Zapater *et al.*, 1986), (ii) from a plasmid for the 5S rDNA probe (pCT4.2; Campell *et al.*, 1992), (iii) from BACs containing the 45S rDNA repeats (BAC T15P10), and (iv) for painting a part of chromosome territory 1 bottom (CT1B) from BACs arranged in contigs (BACs F11P17 to F12B7). The BACs were obtained from the Arabidopsis Biological Resource Center (Ohio, USA). The probes were labeled with modified dUTPs conjugated with Texas-red (Invitrogen) or Alexa-488 (Invitrogen) by nick-translation. The FISH was performed as previously described (Pecinka *et al.*, 2004).

### Indirect immunofluorescence labeling

Nuclei and chromosome preparations were washed in 1×PBS and incubated for 30 min at 37°C in a moist chamber with 30 μl blocking buffer (4% BSA, 0.1% Tween-20 in 1×PBS) to reduce non-specific antibody binding. After three washes in 1×PBS, the slides were incubated with the primary antibodies diluted in antibody buffer (1% BSA, 0.1% Tween-20 in 1×PBS) overnight at 4°C. Next day, the slides were washed in 1×PBS again and incubated with the secondary antibodies in antibody buffer for 1 h at 37°C. After incubation, the preparations were washed in 1×PBS, dehydrated in an ethanol series (70%, 90% and 96% ethanol for 2 min each) and counterstained with DAPI in antifade (Vectashield). All primary and secondary antibodies, and the dilutions used are listed in Table S5.

Immunolocalization of 5-methyl-cytosine requires an initial DNA denaturation of the specimen. Therefore, slides with sorted nuclei were denatured in 70% formamid in 2×SSC for 2 min at 70°C. The preparations were dehydrated in ice cold 70% and 96% ethanol for 5 min each, and air-dried. Subsequent blocking and antibody incubation were carried as described above.

### Microscopy and image analysis

Wide-field fluorescence microscopy was used to evaluate and image the nuclei and chromosome preparations with an Olympus BX61 microscope (Olympus) and an ORCA-ER CCD camera (Hamamatsu). When higher resolution was needed to analyze the spatial arrangement of the chromocenters, a super-resolution fluorescence microscope Elyra PS.1 and the software ZEN (Carl Zeiss GmbH) was used. Processing and analysis of microscopic image stacks were done with ZEN, Adobe Photoshop CS5 (Adobe) and Imaris 8.0 (Bitplane) software.

The CT1B signals were quantified on 16-bit gray scale microscopic images using ImageJ v1.50i (Schneider *et al.*, 2012). The images were taken from preparations of flow-sorted nuclei. Since this technique flattens the nuclei, they were considered as two-dimensional. Within each dataset all images were treated the same way after using the same acquisition parameters. For the CT1B signal image dataset, the background was subtracted with the option ‘Rolling ball’ set at 25 pixels and the delimitation of the region of interest (ROI) with the RenyiEntropy threshold. For the nuclear area image dataset (measured based on DAPI staining), the background was not subtracted, and the nuclear area was delimited as a ROI also with the RenyiEntropy threshold. The area of each ROI was automatically measured by the program.

### Southern blot analysis

Five μg of genomic DNA from *A. thaliana* leaves was digested with either *Hpa*II or *Msp*I (Thermo Fischer Scientific). The DNA was gel-separated and transferred onto a nylon membrane (Hybond XL, Amersham). The ^32^P-labelled centromeric 180 bp repeat pAL was used for Southern hybridization and the signals were detected by autoradiography. The *A. thaliana* centromeric pAL probe was generated by PCR and ^32^P-labeled according to manufacturer’s instructions (Deca-Label DNA labelling Kit, Thermo Scientific).

### *cap-d3* RNA-seq and *in silico* transcriptome analysis

*cap-d3 SAIL*, *cap-d3 SALK* and control (Col-0) seeds were sown in soil and grown under short day conditions. RNA was extracted with the RNeasy Plant Mini Kit (Qiagen) from 50 mg of pooled 4 weeks old plantlets cut above the root. For each of the three *A. thaliana* genotypes five independent RNA extractions were performed and the RNA integrity of the samples was measured in a 2100 Bioanalyzer (Agilent). The four RNA samples of each genotype with the highest RIN (RNA Integrity Number) were used for library preparation and RNA sequencing (NGS platform, IPK Gatersleben, Germany). The libraries were prepared with a TruSeq RNA Library Kit (Illumina) unstranded and sequenced in a HiSeq2000 system (single 100 bp reads).

The quality of the RNA-seq reads were assessed with FastQC v0.11.4 (Babraham Bioinformatics) and adaptors trimmed with Trimmomatic v0.32 (Bolger *et al.*, 2014). After a second quality check in FastQC, the reads were aligned with GSNAP v2016-05-25 (Wu and Nacu, 2010) against the Arabidopsis TAIR10 genome and the gene counts calculated with HTseq v0.6.1 (Anders *et al.*, 2015). Differential expression analyses were performed using the DESeq2 1.14.0 Bioconductor package (Love *et al.*, 2014). Genes were considered differentially expressed (DEG) when they had a Benjamini-Hochberg-adjusted-*P* value ≤ 0.05 and a log_2_-fold change ≤ −1 or ≥ 1. These steps were performed through Galaxy (Afgan *et al.*, 2018).Genes detected as differentially expressed for both *cap-d3* mutants were considered as the genes associated to CAP-D3 defective proteins independently of the specific mutation. Gene enrichment was analyzed with agriGO v1.2 (Du *et al.*, 2010). The analysis of the transcription factors present in *cap-d3* DEG was perform with the Arabidopsis Transcription Factor Database (AtTFDB) from the Arabidopsis Gene Regulatory Information Server (AGRIS; Yilmaz *et al.*, 2011).

### Gene and protein identification numbers

Sequence data from this study can be found in The Arabidopsis Information Resource (TAIR, www.arabidopsis.org) or National Center for Biotechnology Information (NCBI, www.ncbi.nlm.nih.gov/) databases under the following gene identification numbers: *CAP-D2*, AT3G57060; *CAP-D3*, AT4G15890; *CAP-G*, AT5G37630; *CAP-H*, AT2G32590.

## ACKNOWLEDGEMENTS

We thank Jörg Fuchs for flow sorting of nuclei, Katrin Kumke, Oda Weiss, Karla Meier and Ulrike Gresch for excellent technical assistance, Axel Himmelbach for sequencing, and Ingo Schubert for critical reading of the manuscript. The work was supported by a Marie Curie Initial Training Network fellowship (FP7-PEOPLE-2013-ITN, CHIP-ET).

## Author contributions

VS, CM, AH and KG conceived the study and designed the experiments. AH and VS contributed equally to supervise the project. CM, WA, and VS performed the experiments. CM, EK, MVB, IE, FC, AB and IL performed the bioinformatic analysis. CM and VS wrote the manuscript. All authors read and approved the final manuscript.

## Supplementary Data

**Figure S1. *In silico* analysis of *A. thaliana CAP-D2* and *CAP-D3* expression.** The results obtained with the Arabidopsis eFP Browser 2.0 (bar.utoronto.ca) revealed a similar expression level for both genes with high (red), medium (orange) and low (yellow) expression in different organs and developmental stages.

**Figure S2. Protein-protein interaction network of CAP-D2 (condensin I) and CAP-D3 (condensin II).** Both *A. thaliana* CAP-D2 (a, c) and CAP-D3 (b, c) proteins (red) interact potentially with the other coiled-coil condensin SMC complex components (green) and the condensin I- and condensin II-specific subunits (yellow). The network was generated by the STRING program (http://string-db.org/) analysis at scores >0.90 (a, b) and >0.70 (c), respectively. The black lines in between the proteins indicate the supporting evidence from experimental data available from different species. The dashed lines embrace the condensin I and II subunits in (c).

**Figure S3. Affinity purified CAP-D2 and CAP-D3 GS-tagged.** Coomasie staining of a SDS-PAGE gel with protein extracts from cells expressing CAP-D2-GS and CAP-D3-GS. The asterisks indicate the CAP-D2-GS (176 kDa) and CAP-D3-GS (163 kDa) proteins, respectively.

**Figure S4. CAP-G and CAP-H fused to EYFPc localize in the nucleus (arrows) and cytoplasm of *A. thaliana* protoplasts.** The dark regions are chloroplasts.

**Figure S5. CAP-D2, CAP-H and CAP-G fused to EYFPc localize in the nucleus (arrows) and cytoplasm of *N. benthamiana*** leaf epidermal cells

**Figure S6. Western blot analysis confirms the correct size of the CAP-D2 recombinant protein (CAP-D2ct).** Tested against anti-His-tag and anti-T7-tag the recombinant protein produced in *E. coli* has a the expected weight of 59.92 kDa including the T7- and His-tags on the N-t and C-termini, respectively. The arrow marks the band containing the CAP-D2_ct recombinant protein.

**Figure S7. Western blot on different amounts (1–100 ng) of the CAP-D2_ct recombinant protein against the anti-CAP-D2 serum indicates the high sensitivity of anti-CAP-D2.**

**Figure S8. Immunolocalization of histone modifications in *cap-d3* mutants and wild-type plants.** No differences were detected in 4C nuclei of wild-type (Wt) and the *cap-d3 SAIL*, *cap-d3* SALK mutants tested with antibodies against histone H3K27me3 (euchromatic); H3K9me2 (heterochromatic); H3K9ac and with antibodies recognizing H3K14+18+23+27ac.

---

**Table S1.** List of proteins co-purified with CAP-D2-GS. Only proteins present in the three affinity purifications and with no occurrence among the non-specific proteins are listed.

| AGI | Protein name/function | Average MASCOT Score |
| --- | --- | --- |
| AT3G57060 | CAP-D2 | 4538.87 |
| AT5G48600 | SMC4 | 2542.57 |
| AT5G62410 | SMC2A | 2497.43 |
| AT3G47460 | SMC2B | 1932.23 |
| AT5G37630 | CAP-G | 1421.20 |
| AT2G32590 | CAP-H | 919.63 |
| AT3G46740 | protein TOC75-3 | 641.57 |
| AT3G08943 | armadillo/beta-catenin-like repeat-containing protein | 551.70 |
| AT1G72560 | PAUSED | 476.33 |
| AT5G09840 | putative endonuclease or glycosyl hydrolase | 470.07 |
| AT2G20800 | NAD(P)H dehydrogenase B4 | 445.47 |
| AT2G02090 | CHR19 Chromatin remodelling 19 | 400.83 |
| AT5G64270 | putative splicing factor | 393.70 |
| AT4G24840 | Brefeldin A-sensitive Golgi protein-like | 389.23 |
| AT1G07810 | Ca<sup>2+</sup>-transporting ATPase | 386.87 |
| AT4G01100 | adenine nucleotide transporter 1 | 380.70 |
| AT3G60860 | guanine nucleotide exchange factor | 374.00 |
| AT4G19490 | protein VPS54 | 360.83 |
| AT3G54110 | uncoupling mitochondrial protein 1 | 352.93 |
| AT2G40730 | SCY1-like protein | 352.83 |
| AT4G02510 | translocase of chloroplast 159 | 341.17 |
| AT3G01280 | mitochondrial outer membrane protein porin 1 | 340.97 |
| AT4G05020 | NAD(P)H dehydrogenase B2 | 334.20 |
| AT5G41950 | tetratricopeptide repeat domain-containing protein | 332.53 |
| AT5G16930 | AAA-type ATPase family protein | 329.23 |
| AT4G01400 | oligomeric golgi complex subunit-like protein | 318.60 |
| AT4G02570 | CUL1 cullin 1 | 315.07 |
| AT5G16210 | HEAT repeat-containing protein | 301.20 |
| AT2G39260 | regulator of nonsense transcripts UPF2 | 269.63 |
| AT5G22770 | AP-2 complex subunit alpha-1 | 265.63 |
| AT5G18420 | CCR4-NOT transcription complex subunit | 265.13 |
| AT2G01690 | ARM repeat superfamily protein | 256.73 |
| AT3G45190 | SIT4 phosphatase-associated family protein | 253.40 |
| AT4G33650 | dynamamin like protein 2a | 250.40 |
| AT4G02350 | exocyst complex component sec15B | 247.90 |
| AT2G36200 | kinesin family member 11 | 246.07 |
| AT5G26760 | RPAP2 IYO MATE (RIMA) | 243.80 |
| AT1G04080 | pre-mRNA-processing factor 39 | 242.80 |
| AT3G62360 | carbohydrate-binding-like fold-containing protein | 241.90 |
| AT1G60200 | RNA-Binding protein 25 | 236.77 |
| AT4G32050 | neurochondrin family protein | 234.23 |
| AT1G22730 | putative topoisomerase | 226.80 |
| AT5G13850 | nascent polypeptide-associated complex subunit alpha-like protein 3 | 226.57 |
| AT3G55410 | 2-oxoglutarate dehydrogenase, E1 subunit-like protein | 218.97 |
| AT2G18330 | AAA-type ATPase-like protein | 218.27 |
| AT3G16830 | Topless-related 2 protein | 218.23 |
| AT2G22300 | calmodulin-binding transcription activator 3 | 213.63 |
| AT2G14120 | dynamamin-like protein | 204.30 |
| AT4G01990 | tetratricopeptide repeat-like superfamily protein | 203.83 |
| AT3G11710 | lysyl-tRNA synthetase | 202.50 |
| AT5G49830 | exocyst complex component 84B | 197.60 |
| AT2G27170 | SMC3 | 196.17 |
| AT1G48900 | signal recognition particle subunit SRP54 | 195.77 |
| AT1G60070 | adaptor protein complex AP-1, gamma subunit | 194.00 |
| AT2G27900 | coiled-coil protein | 193.10 |
| AT1G63810 | nucleolar protein | 192.43 |
| AT4G21150 | ribophorin II (RPN2) family protein | 187.57 |
| AT2G31810 | ACT domain-containing small subunit of acetolactate synthase protein | 186.10 |
| AT4G27500 | proton pump interactor 1 | 186.07 |
| AT1G73430 | putative conserved oligomeric golgi complex 3 | 182.73 |
| AT1G71270 | A. thaliana VPS52 homolog | 178.03 |
| AT5G19760 | mitochondrial substrate carrier family protein | 176.33 |
| AT3G54540 | ABC transporter F family member 4 | 175.57 |
| AT5G13110 | glucose-6-phosphate 1-dehydrogenase | 171.50 |
| AT5G08550 | GC-rich sequence DNA-binding factor | 167.73 |
| AT5G47480 | RGPR-related protein | 167.73 |
| AT5G65460 | kinesin like protein for actin based chloroplast movement 2 | 167.23 |
| AT5G14580 | polyribonucleotide nucleotidyltransferase | 163.70 |
| AT5G09420 | translocon at the outer membrane of chloroplasts 64-V | 162.17 |
| AT3G11400 | eukaryotic translation initiation factor 3g | 155.40 |
| AT5G08450 | HDC1 histone deacetylation complex 1 | 153.20 |
| AT5G11980 | conserved oligomeric Golgi complex component-related | 153.10 |
| AT1G79940 | translocation protein SEC63 | 152.73 |
| AT3G45970 | expansin-like A1 | 152.73 |
| AT2G42710 | ribosomal protein .1/L10 family protein | 147.03 |
| AT3G08030 | uncharacterized protein | 146.87 |
| AT5G18620 | CHR17 chromatin remodeling factor17 | 146.83 |
| AT5G50320 | ELO3 histone acetyltransferase | 146.53 |
| AT1G31780 | conserved oligomeric Golgi complex 7 | 144.90 |
| AT5G19400 | telomerase activating protein Est1 | 142.17 |
| AT5G10470 | geminivirus Rep-interacting motor protein | 138.43 |
| AT1G61040 | Plus-3 domain-containing protein | 137.57 |
| AT1G67930 | Golgi transport complex-related protein | 136.70 |
| AT4G39690 | homolog of yeasst mic60 protein | 136.10 |
| AT1G03860 | prohibitin 2 | 135.43 |
| AT1G19870 | protein IQ-domain 32 | 135.23 |
| AT3G16620 | translocase of chloroplast 120 | 133.90 |
| AT5G61970 | signal recognition particle subunit SRP68 | 133.60 |
| AT1G32380 | phosphoribosyl pyrophosphate synthetase II | 128.47 |
| AT1G78380 | glutathione S-transferase TAU 19 | 128.00 |
| AT3G59020 | armadillo/beta-catenin-like repeat-containing protein | 125.70 |
| AT4G10320 | similar to isoleucyl-tRNA synthetases | 121.20 |
| AT4G24550 | AP-4 complex subunit mu-1 | 121.07 |
| AT1G62740 | putative stress-inducible protein | 117.43 |
| AT1G11910 | aspartic proteinase | 117.33 |
| AT5G67500 | voltage dependent anion channel 2 | 116.70 |
| AT5G46750 | ARF-GAP domain 9 | 115.70 |
| AT2G19480 | putative nucleosome assembly protein 1;2 | 115.53 |
| AT5G66680 | putative dolichyl-di-phosphooligosaccharide-protein glycotransferase | 114.07 |
| AT5G03540 | exocyst subunit exo70 family protein A1 | 112.07 |
| AT5G42960 | Outer envelope pore 24B-like protein | 110.17 |
| AT3G46220 | E3 UFM1-protein ligase 1-like protein | 109.93 |
| AT2G27030 | calmodulin 5 | 109.70 |
| AT3G44330 | M28 Zn-peptidase nicastrin | 109.40 |
| AT3G23300 | S-adenosyl-Lmethionine-dependent methyltransferases superfamily protein | 107.67 |
| AT1G49040 | stomatal cytokinesis defective (SCD1) | 106.07 |
| AT2G05120 | nucleoporin, Nup133/Nup155-like protein | 103.40 |

**Table S2.** List of proteins co-purified with CAP-D3-GS. Only proteins present in the three affinity purifications and with no occurrence among the non-specific proteins are listed.

| AGI | Protein names/function | Average MASCOT Score |
| --- | --- | --- |
| AT4G15890 | CAP-D3 | 6668.17 |
| AT5G48600 | SMC4 | 1571.70 |
| AT5G62410 | SMC2A | 819.37 |
| AT2G19480 | nucleosome assembly protein 1;2 | 496.57 |
| AT2G38770 | intron-binding protein aquarius | 440.83 |
| AT3G13290 | varicose-related protein | 416.07 |
| AT4G02510 | translocase of chloroplast 159 | 407.33 |
| AT3G54110 | uncoupling mitochondrial protein 1 | 399.83 |
| AT1G48900 | signal recognition particle subunit SRP54 | 399.30 |
| AT4G26110 | nucleosome assembly protein 1-like 1 | 397.30 |
| AT4G01990 | pentatricopeptide repeat-containing protein | 371.23 |
| AT4G32050 | Neurochondrin family protein | 331.57 |
| AT2G03510 | SPFH/Band 7/PHB domain-containing membrane-associated protein | 317.63 |
| AT4G01100 | adenine nucleotide transporter 1 | 309.07 |
| AT3G02200 | proteasome component (PCI) domain protein | 308.57 |
| AT2G42710 | ribosomal protein .1/L10 family protein | 304.23 |
| AT3G60860 | SEC7-like guanine nucleotide exchange family protein | 303.10 |
| AT3G01280 | mitochondrial outer membrane protein porin 1 | 299.57 |
| AT5G09840 | putative endonuclease or glycosyl hydrolase | 288.67 |
| AT4G21150 | ribophorin II (RPN2) family protein | 250.40 |
| AT5G18420 | CCR4-NOT transcription complex subunit | 245.07 |
| AT1G64960 | CAP-G2 | 218.27 |
| AT4G33510 | phospho-2-dehydro-3-deoxyheptonate aldolase 2 | 211.13 |
| AT5G58410 | HEAT/U-box domain-containing protein | 210.40 |
| AT5G19760 | mitochondrial substrate carrier family protein | 207.50 |
| AT3G16730 | CAP-H2 | 205.97 |
| AT1G14850 | nucleoporin 155 | 200.63 |
| AT1G06530 | tropomyosin-related protein | 198.07 |
| AT5G40770 | prohibitin 3 | 194.70 |
| AT3G02650 | pentatricopeptide repeat-containing protein | 186.60 |
| AT2G20800 | NAD(P)H dehydrogenase B4 | 184.30 |
| AT1G20960 | putative U5 small nuclear ribonucleoprotein helicase | 183.03 |
| AT2G45140 | VAP-like protein 12 | 182.27 |
| AT3G44330 | M28 Zn-peptidase nicastrin | 180.40 |
| AT5G13110 | glucose-6-phosphate 1-dehydrogenase | 177.80 |
| AT4G24550 | AP-4 complex subunit mu-1 | 174.97 |
| AT5G64270 | putative splicing factor | 170.03 |
| AT5G66680 | putative dolichyl-di-phosphooligosaccharide-protein glycotransferase | 168.17 |
| AT3G02090 | putative mitochondrial processing peptidase | 162.33 |
| AT2G39260 | regulator of nonsense transcripts UPF2 | 161.90 |
| AT5G07340 | calnexin homolog | 155.57 |
| AT2G33040 | ATP synthase subunit gamma | 155.37 |
| AT4G02150 | Importin subunit alpha-2 | 155.03 |
| AT5G50320 | ELO3 histone acetyl transferase | 154.50 |
| AT3G55620 | translation initiation factor IF6 | 153.50 |
| AT5G27970 | armadillo/beta-catenin-like repeat-containing protein | 153.37 |
| AT1G26460 | pentatricopeptide repeat-containing protein | 153.33 |
| AT1G55890 | pentatricopeptide repeat-containing protein | 152.80 |
| AT1G02370 | pentatricopeptide repeat-containing protein | 151.40 |
| AT5G11980 | putative CONSERVED OLIGOMERIC GOLGI COMPLEX 8 | 148.23 |
| AT5G15610 | proteasome component (PCI) domain protein | 146.13 |
| AT5G20490 | Myosin XI family protein with Dil domain | 146.00 |
| AT5G30510 | ribosomal protein S1 | 145.93 |
| AT4G17330 | hypothetical protein | 144.13 |
| AT5G15020 | SNL2 homolog of the transcriptional repressor SIN3 | 142.40 |
| AT2G20360 | NAD(P)-binding Rossmann-fold superfamily protein | 141.63 |
| AT2G31810 | ACETOLACTATE SYNTHASE SMALL SUBUNIT 1 | 140.57 |
| AT1G64880 | hypothetical Protein | 139.73 |
| AT5G16930 | AAA-type ATPase family protein | 134.10 |
| AT1G27090 | uncharacterized glycine-rich protein | 132.70 |
| AT1G71410 | SCYL2B | 132.50 |
| AT3G55005 | TONNEAU 1B. Involved in cortical microtubule organization | 129.97 |
| AT3G53130 | carotenoid epsilon-ring hydroxylase | 129.13 |
| AT3G49080 | RIBOSOMAL PROTEIN S9 M | 128.83 |
| AT2G43950 | chloroplast outer envelope protein 37 | 128.77 |
| AT2G26890 | GRAVITROPISM DEFECTIVE 2 | 126.27 |
| AT5G12470 | RER4 putative UvrABC system C protein | 125.97 |
| AT4G31810 | 3-hydroxyisobutyryl-CoA hydrolase-like protein 2 | 122.43 |
| AT3G46950 | mitochondrial transcription termination factor family protein | 119.33 |
| AT3G18790 | pre-mRNA-splicing factor ISY1 | 116.20 |
| AT1G23280 | MAK16 protein-like protein | 114.93 |
| AT1G67140 | HEAT repeat-containing protein | 113.27 |
| AT5G46750 | putative ADP-ribosylation factor GTPase-activating protein AGD9 | 110.80 |
| AT3G61140 | CSN1 COP9 signalosome 1 | 110.77 |
| AT2G46020 | ATP-dependent helicase BRAHMA | 109.03 |
| AT4G21800 | QQT2. Required for early embryo development. | 108.77 |
| AT2G15630 | pentatricopeptide repeat-containing protein | 108.30 |
| AT4G38600 | E3 ubiquitin-protein ligase UPL3 | 108.17 |
| AT2G40890 | putative cytochrome P450 | 103.80 |
| AT4G11260 | phosphatase SGT1b | 103.67 |
| AT1G50030 | FKBP12-rapamycin complex-associated protein | 102.90 |
| AT4G05020 | NAD(P)H dehydrogenase B2 | 101.63 |
| AT2G16485 | NERD (Needed for RDR2-independent DNA methylation) | 101.37 |
| AT2G16640 | putative chloroplast outer membrane protein | 100.20 |

**Table S3.**
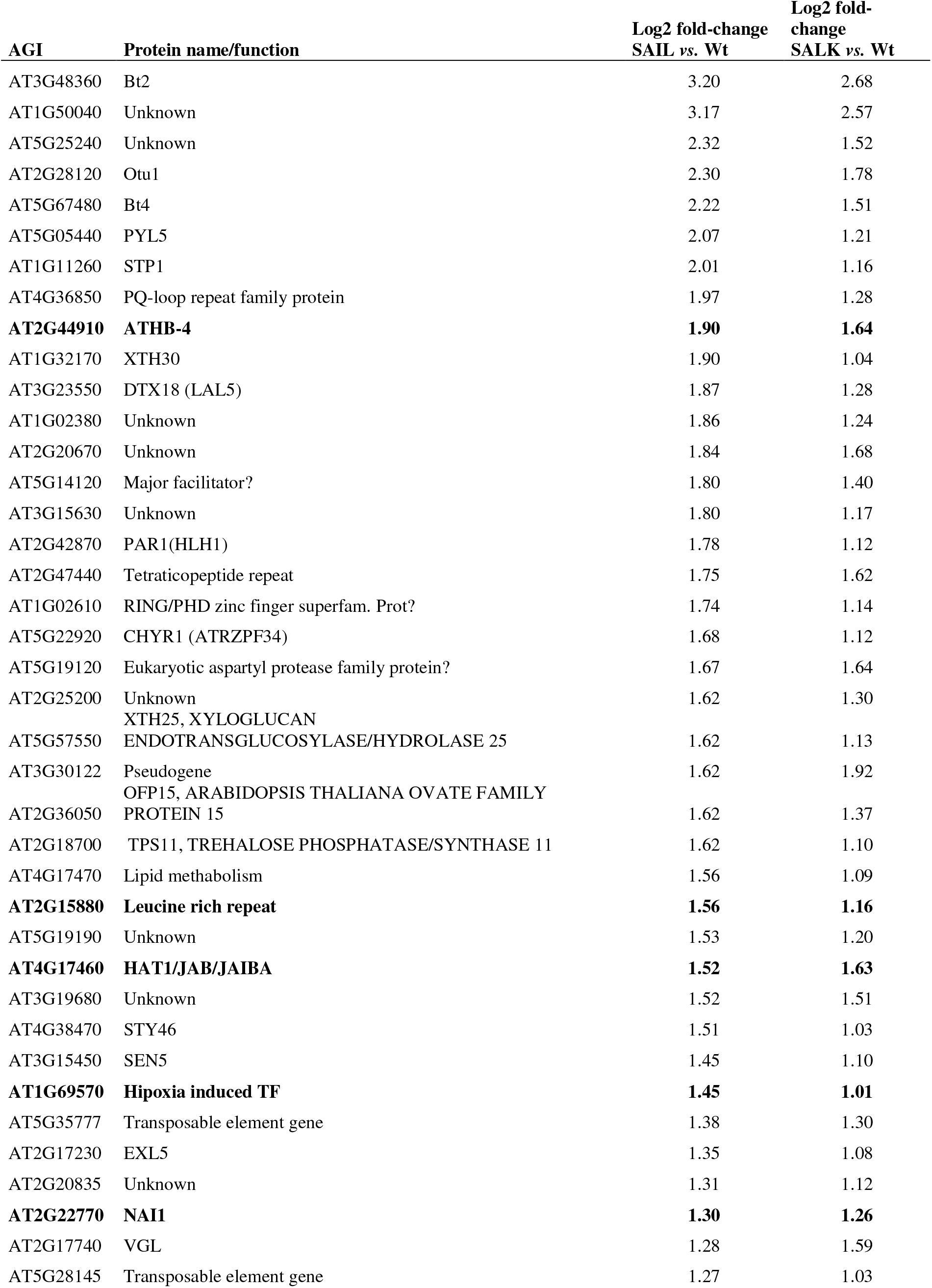
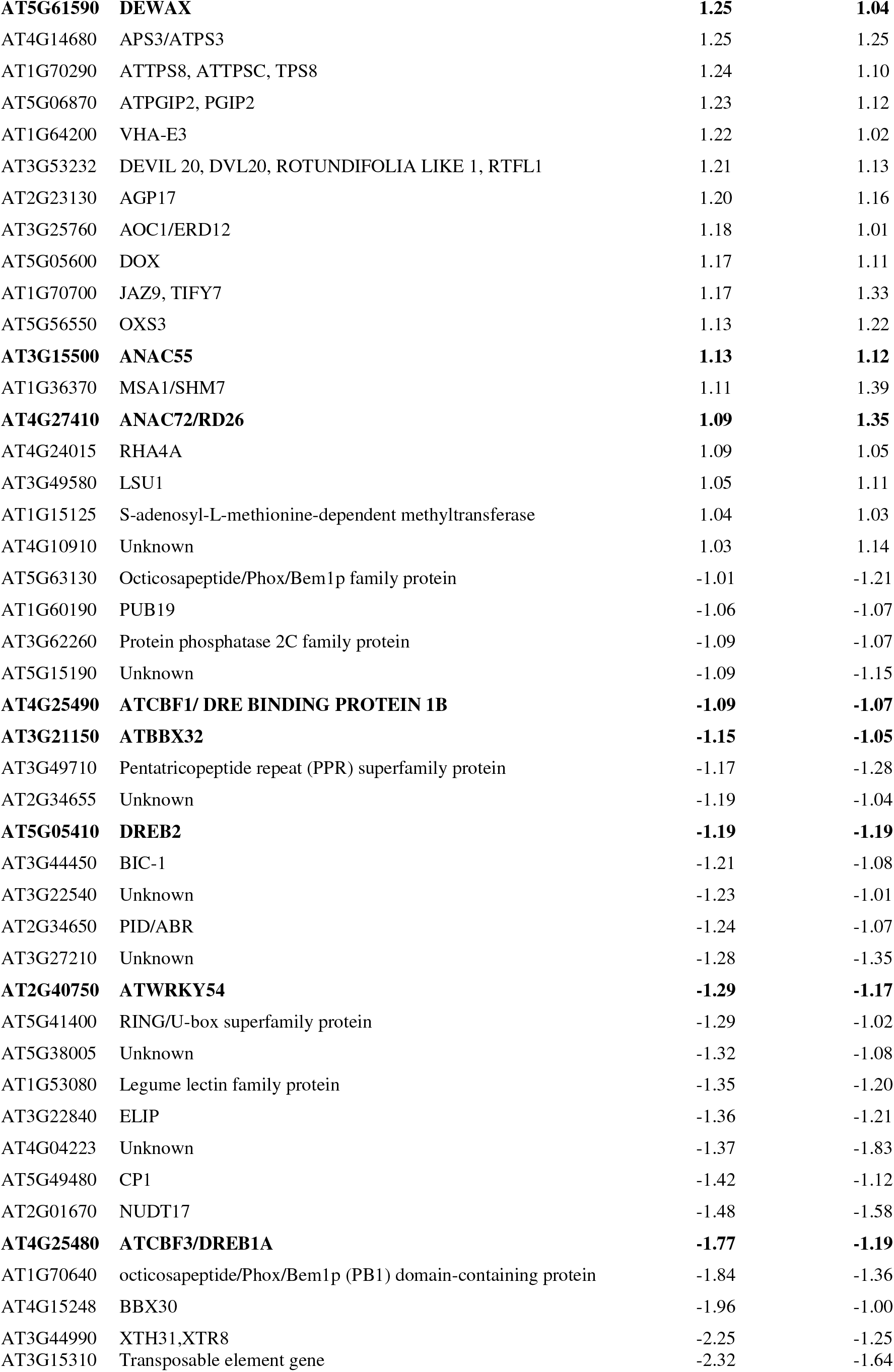
Differentially expressed genes (DEGs) in the *cap-d3* mutants. List of DEGs common to *cap-d3 SAIL vs.* wild-type (Wt) and *cap-d3 SALK vs*. Wt. Genes in bold are transcription factors.

**Table S4.**
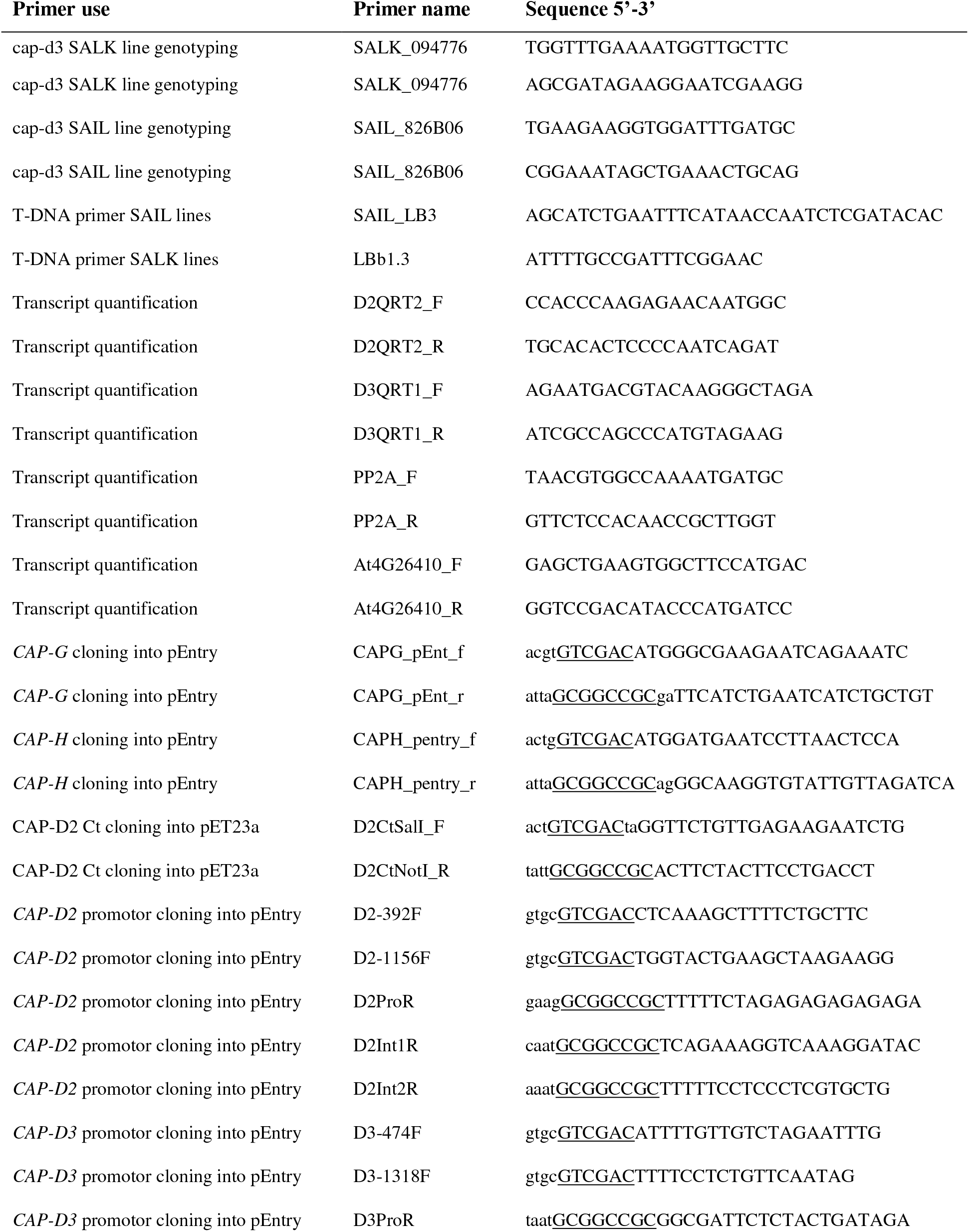
Primer sequences and usage. Linker sequences are in lower case letters, restriction sites are underlined and genomic sequences are written in upper case.

**Table S5.**
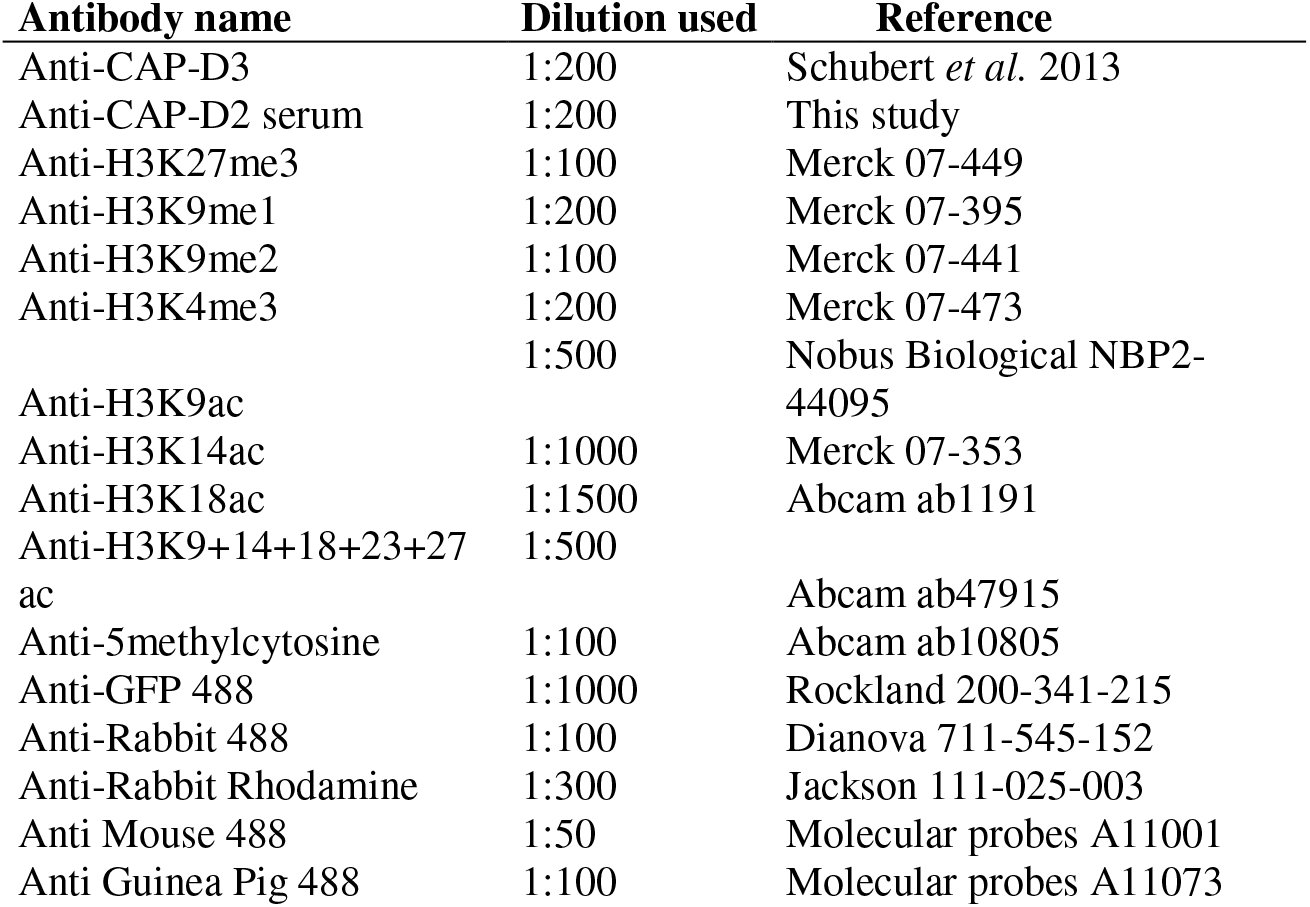
Antibodies and their dilutions used for immunolocalization.

**Figure S1.**
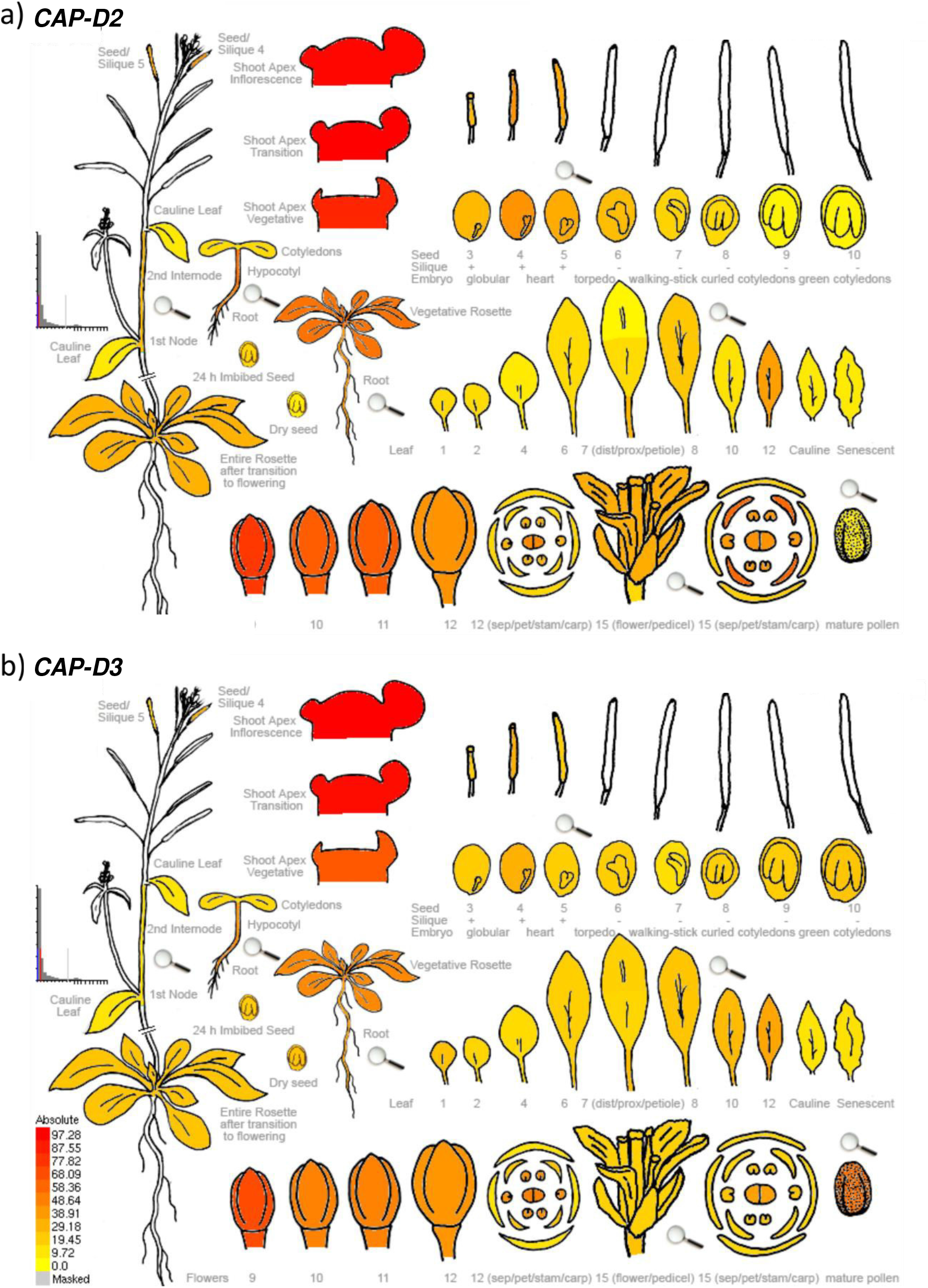
*In silico* analysis of *A. thaliana CAP-D2* and *CAP-D3* expression. The results obtained with the *Arabidopsis* eFP Browser 2.0 (bar.utoronto.ca) revealed a similar expression level for both genes with high (red), medium (orange) and low (yellow) expression in different organs and developmental stages.

**Figure S2.**
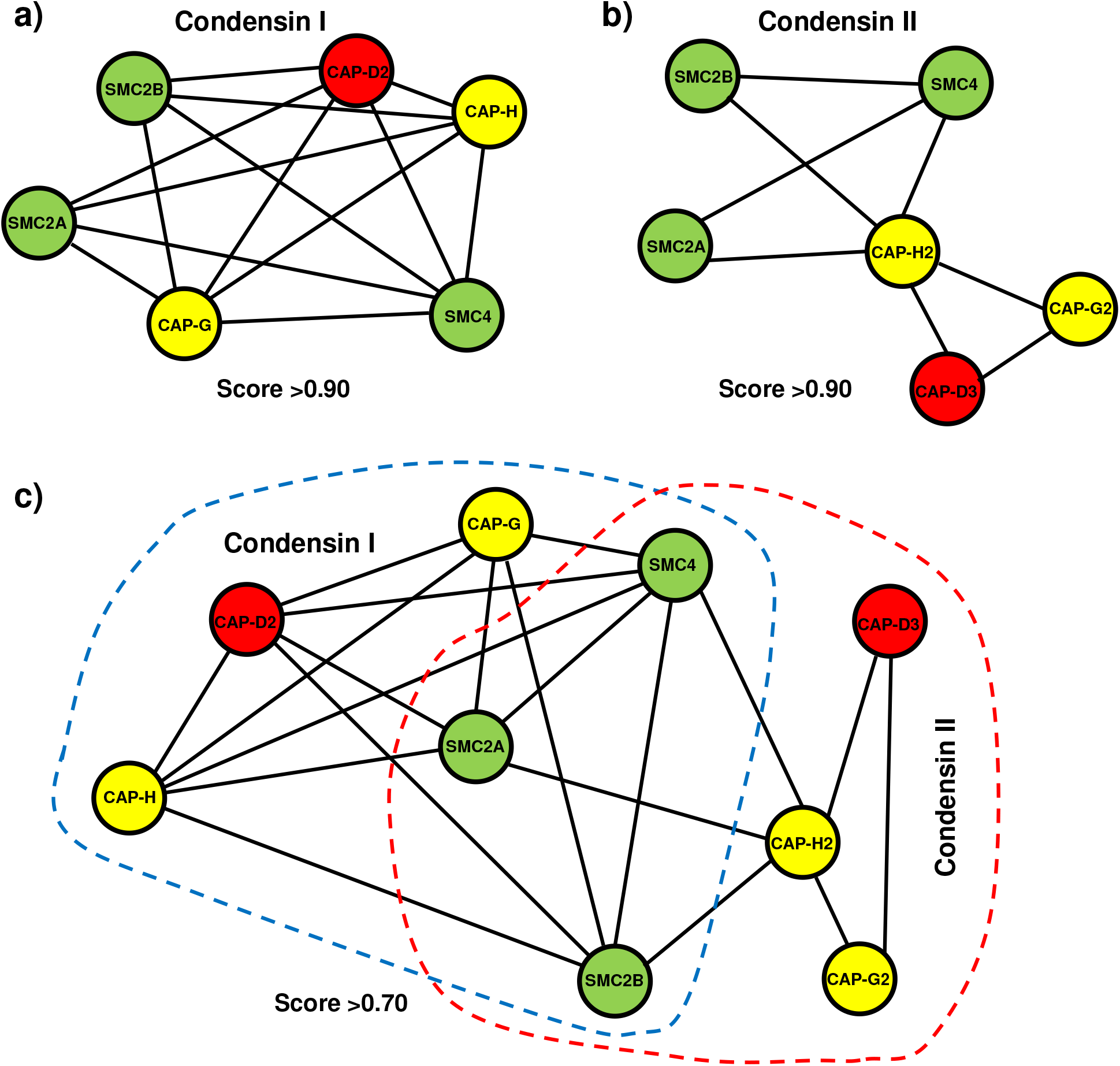
Protein-protein interaction network of CAP-D2 (condensin I) and CAP-D3 (condensin II). Both *A. thaliana* CAP-D2 (a, c) and CAP-D3 (b, c) proteins (red) interact potentially with the other coiled-coil condensin SMC complex components (green) and the condensin I- and condensin II-specific subunits (yellow). The network was generated by a STRING program (http://string-db.org/) analysis at scores >0.90 (a, b) and >0.70 (c), respectively. The black lines in between the proteins indicate the supporting evidence from experimental data available from different species. The dashed lines embrace the condensin I and II subunits in (c).

**Figure S3.**
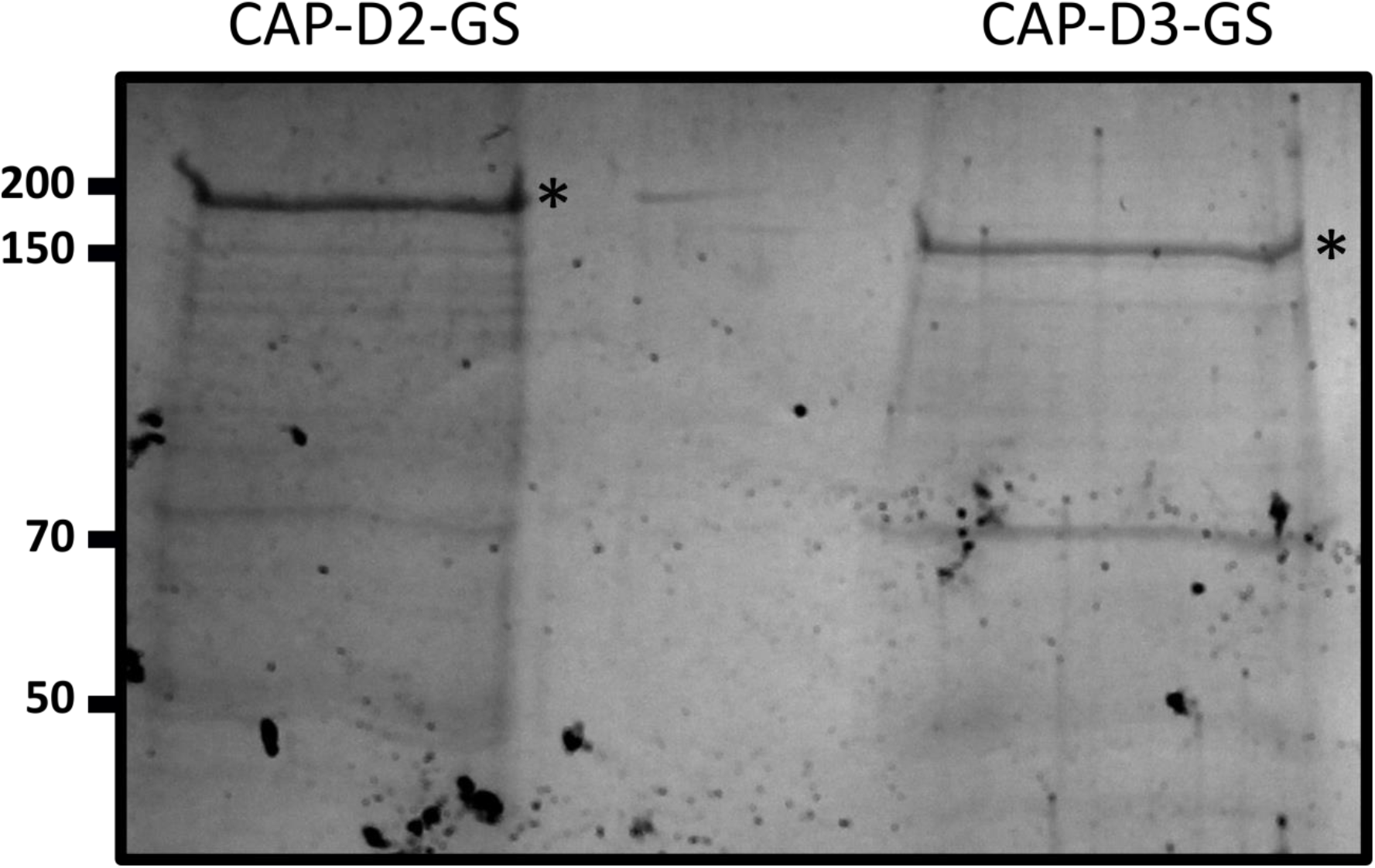
Affinity purified CAP-D2 and CAP-D3 GS-tagged. Coomasie staining of a SDS-PAGE gel with protein extracts from cells expressing CAP-D2-GS and CAP-D3-GS. The asterisks indicate .the CAP-D2-GS (176 kDa) and CAP-D3-GS (163 kDa) proteins, respectively.

**Figure S4.**
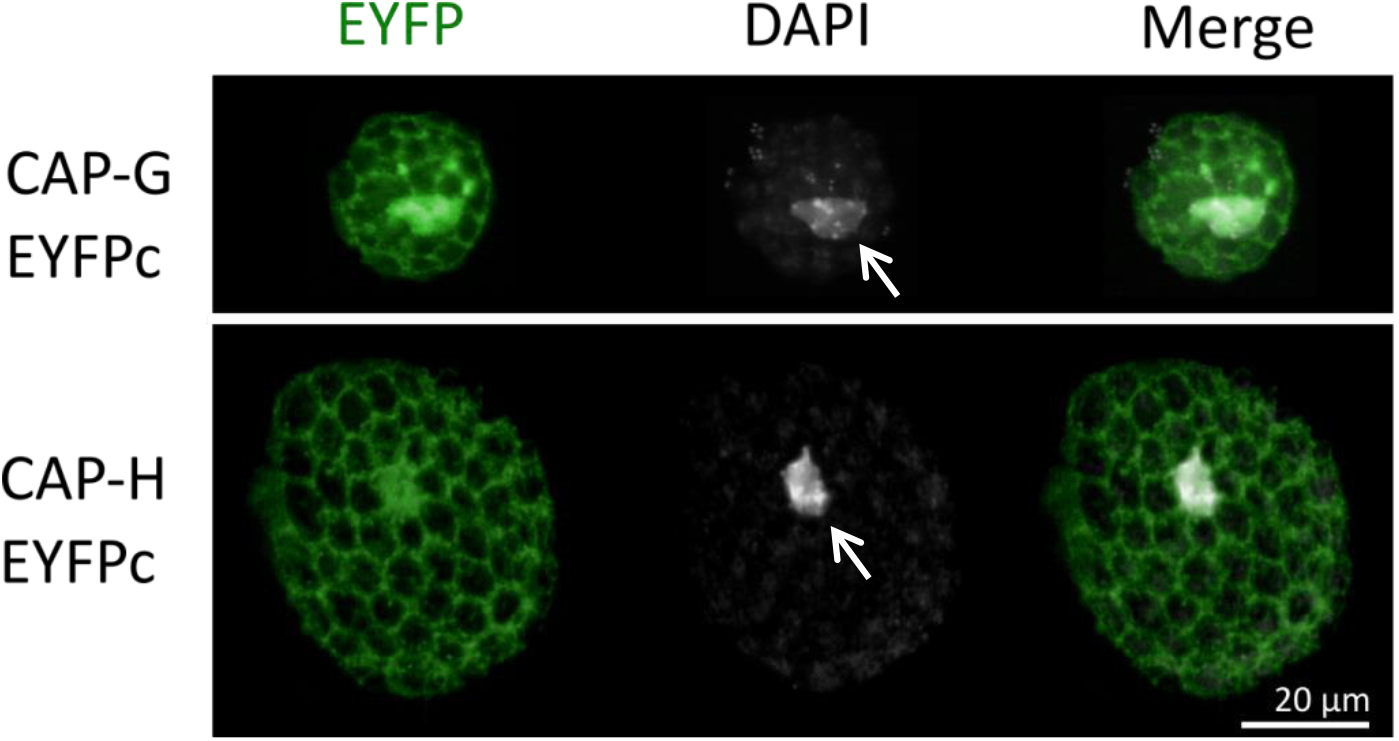
CAP-G and CAP-H fused to EYFPc localize in the nucleus (arrows) and cytoplasm of *A. thaliana* protoplasts. The dark regions are chloroplasts.

**Figure S5.**
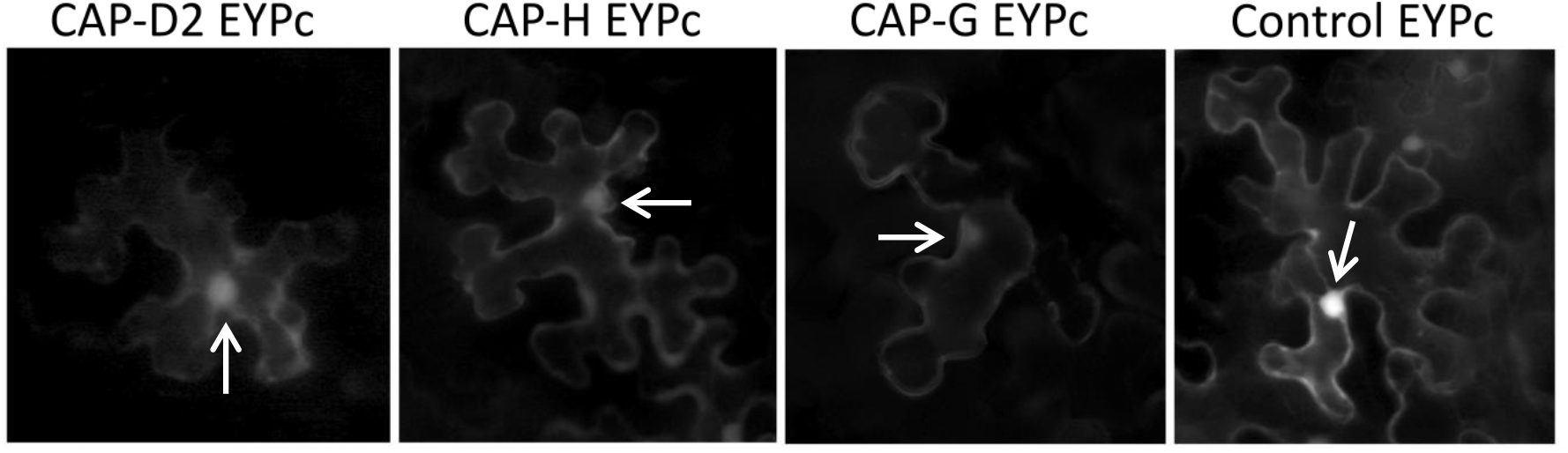
CAP-D2, CAP-H and CAP-G fused to EYFPc localize in the nucleus (arrows) and cytoplasm of *N. benthamiana* leaf epidermal cells.

**Figure S6.**
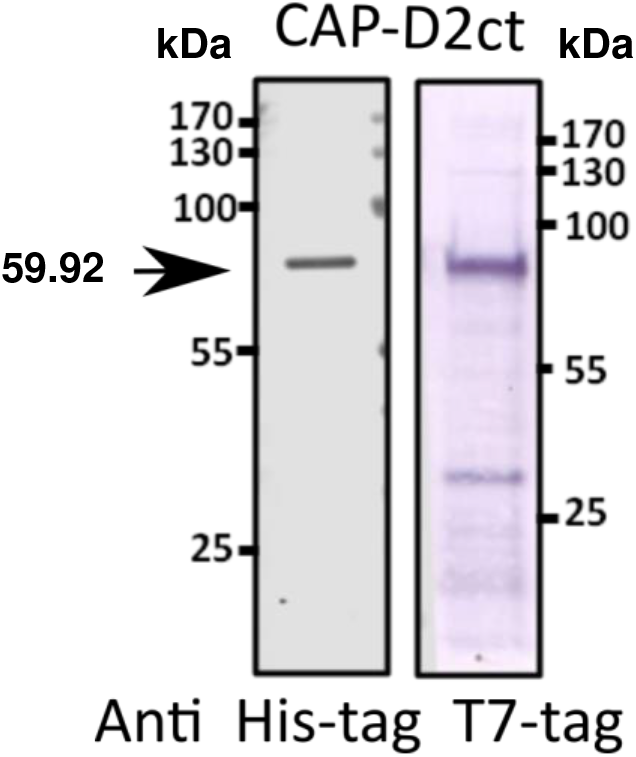
Western blot analysis confirms the correct size of the CAP-D2 recombinant protein (CAP-D2ct). Tested against anti-His-tag and anti-T7-tag the recombinant protein produced in *E. coli* has a the expected weight of 59.92 kDa including the T7- and His-tags on the N-t and C-termini, respectively. The arrow marks the band containing the CAP-D2_ct recombinant protein.

**Figure S7.**
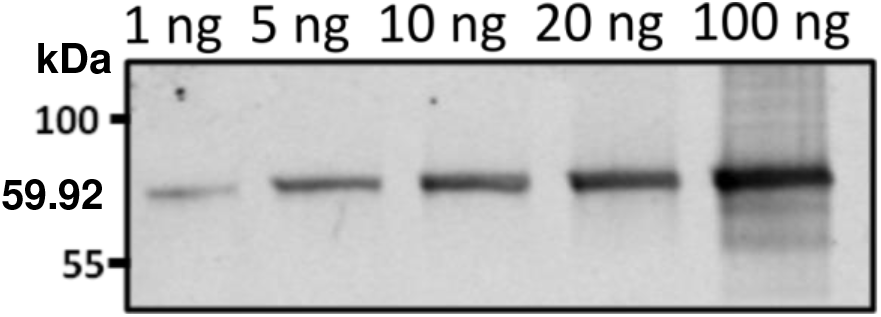
Western blot on different amounts (1–100 ng) of the CAP-D2_ct recombinant protein against the anti-CAP-D2 serum indicates the high sensitivity of anti-CAP-D2.

**Figure S8.**
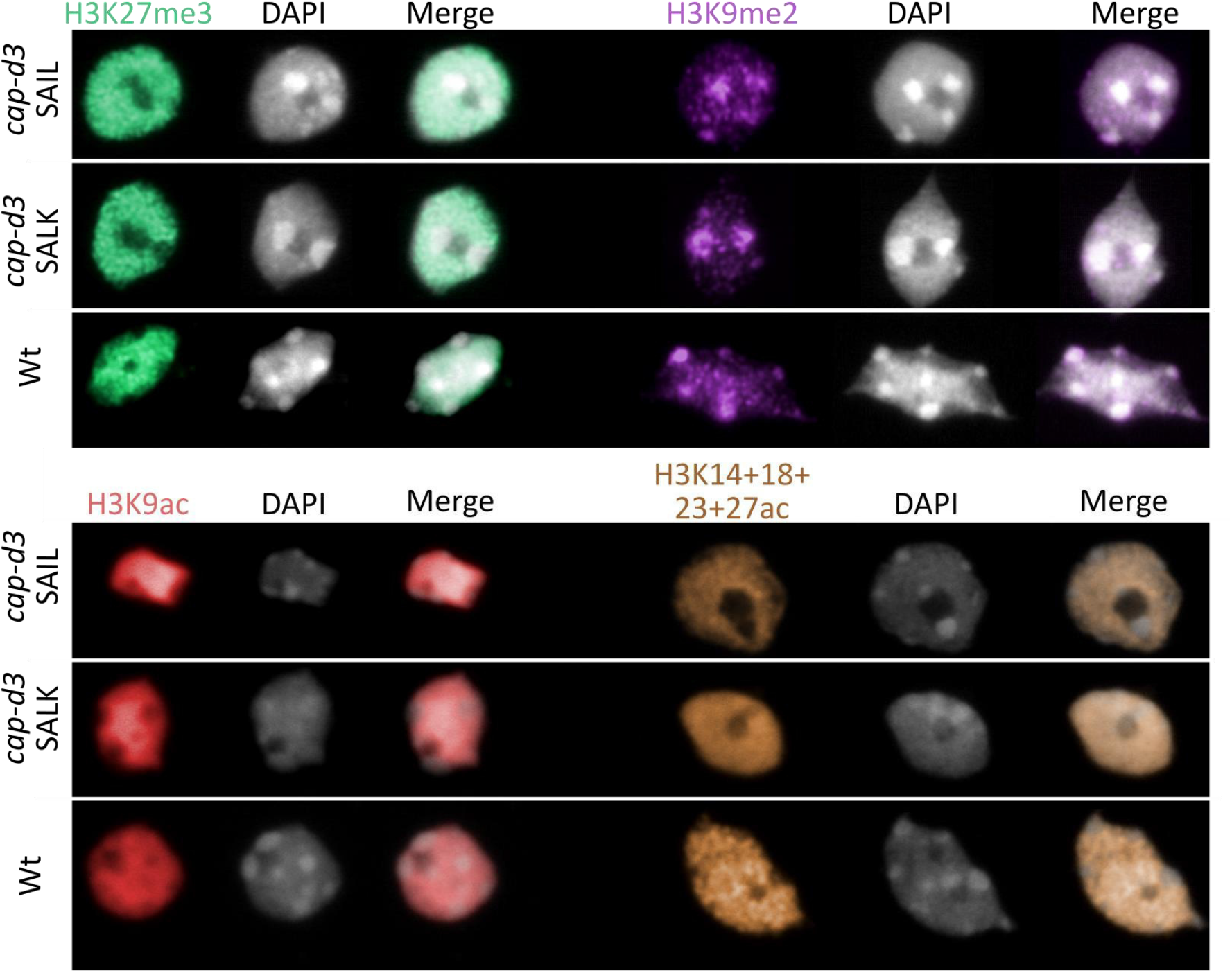
Immunolocalization of histone modifications in *cap-d3* mutants and wild-type plants. No differences were detected in 4C nuclei of wild-type (Wt) and the *cap-d3 SAIL*, *cap-d3 SALK* mutants tested with antibodies against histone H3K27me3 (euchromatic); H3K9me2 (heterochromatic); H3K9ac and with antibodies recognizing H3K14+18+23+27ac.

## REFERENCES

Afgan, E., Baker, D., Batut, B., Van Den Beek, M., Bouvier, D., Čech, M., Chilton, J., Clements, D., Coraor, N., Grüning, B.A. and Guerler, A. (2018) The Galaxy platform for accessible, reproducible and collaborative biomedical analyses: 2018 update. Nucleic Acids Res., 46, W537–W544.

Anders, S., Pyl, P.T. and Huber, W. (2015) HTSeq-a Python frame work to work with high-throughtput sequencing data. Bioinformatics, 31, 166–169.

Andrey, P., Kieu, K., Kress, C., Lehmann, G., Tirichine, L., Liu, Z.C., Biot, E., Adenot, P.G., Hue-Beauvais, C., Houba-Herin, N., Duranthon, V., Devinoy, E., Beaujean, N., Gaudin, V., Maurin, Y. and Debey, P. (2010) Statistical analysis of 3D images detects regular spatial distributions of centromeres and chromocenters in animal and plant nuclei. PloS Computat. Biol., 6, e1000853.

Antosz, W., Pfab, A., Ehrnsberger, H.F., Holzinger, P., Kollen, K., Mortensen, S.A., Bruckmann, A., Schubert, T., Langst, G., Griesenbeck, J., Schubert, V., Grasser, M. and Grasser, K.D. (2017) The composition of the Arabidopsis RNA polymerase II transcript elongation complex reveals the interplay between elongation and mRNA processing factors. Plant Cell, 29, 854–870.

Bauer, C.R., Hartl, T.A. and Bosco, G. (2012) Condensin II promotes the formation of chromosome territories by inducing axial compaction of polyploid interphase chromosomes. PLoS Genet., 8, e1002873.

Berr, A. and Schubert, I. (2007) Interphase chromosome arrangement in *Arabidopsis thaliana* is similar in differentiated and meristematic tissues and shows a transient mirror symmetry after nuclear division. Genetics, 176, 853–863.

Biggs, R., Liu, P.Z., Stephens, A.D. and Marko, J.F. (2019) Effects of altering histone posttranslational modifications on mitotic chromosome structure and mechanics. Mol. Biol. Cell, 30, 820–827.

Bolger, A.M., Lohse, M. and Usadel, B. (2014) Trimmomatic: a flexible trimmer for Illumina sequence data. Bioinformatics, 30, 2114–2120.

Bourbousse, C., Mestiri, I., Zabulon, G., Bourge, M., Formiggini, F., Koini, M.A., Brown, S.C., Fransz, P., Bowler, C. and Barneche, F. (2015) Light signaling controls nuclear architecture reorganization during seedling establishment. Proc. Natl. Acad. Sci. U S A, 112, E2836–2844.

Boyle, S., Gilchrist, S., Bridger, J.M., Mahy, N.L., Ellis, J.A. and Bickmore, W.A. (2001) The spatial organization of human chromosomes within the nuclei of normal and emerin-mutant cells. Hum. Mol. Genet., 10, 211–219.

Buster, D.W., Daniel, S.G., Nguyen, H.Q., Windler, S.L., Skwarek, L.C., Peterson, M., Roberts, M., Meserve, J.H., Hartl, T., Klebba, J.E., Bilder, D., Bosco, G. and Rogers, G.C. (2013) SCFSlimbubiquitin ligase suppresses condensin II-mediated nuclear reorganization by degrading Cap-H2. J. Cell Biol., 201, 49–63.

Campell, B.R., Song, Y., Posch, T.E., Cullis, C.A. and Town, C.D. (1992) Sequence and organization of 5S ribosomal RNA-encoding genes of *Arabidopsis thaliana*. Gene, 112, 225–228.

Chalut, K.J., Hopfler, M., Lautenschlager, F., Boyde, L., Chan, C.J., Ekpenyong A., Martinez-Arias, A. and Guck, J. (2012) Chromatin decondensation and nuclear softening accompany Nanog downregulation in embryonic stem cells. Biophys. J., 103, 2060–2070.

Clough, S.J. and Bent, A.F. (1998) Floral dip: A simplified method for Agrobacterium-mediated transformation of *Arabidopsis thaliana*. Plant J., 16, 735–743.

Cobbe, N. and Heck, M.M.S.(2004) The evolution of SMC proteins: phylogenetic analysis and structural implications. Mol. Biol. Evol., 21, 332–347.

Cobbe, N., Savvidou, E. and Heck, M.M.S. (2006) Diverse mitotic and interphase functions of condensins in Drosophila. Genetics, 172, 991–1008.

Cremer, T. and Cremer, M. (2010) Chromosome territories. Cold Spring Harbor Perspectives in Biology, 2, a003889.

Cremer, T., Cremer, M. and Cremer, C. (2018) The 4D nucleome: genome compartmentalization in an evolutionary context. Biochemistry (Mosc), 83, 313–325.

Damodaran, K., Venkatachalapathy, S., Alisafaei, F., Radhakrishnan, A.V., Sharma Jokhun, D., Shenoy, V.B. and Shivashankar, G.V. (2018) Compressive force induces reversible chromatin condensation and cell geometry dependent transcriptional response. Mol. Biol. Cell, 29, 3039–3051.

De Nooijer, S., Wellink, J., Mulder, B. and Bisseling, T. (2009) Non-specific interactions are sufficient to explain the position of heterochromatic chromocenters and nucleoli in interphase nuclei. Nucleic Acids Res., 37, 3558–3568.

Doğan, E.S. and Liu, C. (2018) Three-dimensional chromatin packing and positioning of plant genomes. Nat. Plants, 8,521–529.

Dowen, J.M., Bilodeau, S., Orlando, D.A., Hübner, M.R., Abraham, B.J., Spector, D.L. and Young, R.A. (2013) Multiple structural maintenance of chromosome complexes at transcriptional regulatory elements. Stem Cell Reports, 1, 371–378.

Du, Z., Zhou, X., Ling, Y., Zhang, Z. and Su, Z. (2010) agriGO: a GO analysis toolkit for the agricultural community. Nucleic Acids Res., 38, W64–70.

Dürr, J., Lolas, I.B., Sørensen, B.B., Schubert, V., Houben, A., Melzer, M., Deutzmann, R., Grasser, M. and Grasser, K.D. (2014) The transcript elongation factor SPT4/SPT5 is involved in auxin-related gene expression in Arabidopsis. Nucleic Acids Res., 42, 4332–4347.

Elbatsh, A.M.O., Kim, E., Eeftens, J.M., Raaijmakers, J.A., van der Weide, R.H., Garcia-Nieto, A., Bravo, S., Ganji, M., Uit de Bos, J., Teunissen, H., Medema, R.H., de Wit, E., Haering, C.H., Dekker, C. and Rowland, B.D. (2019) Distinct roles for condensin’s two ATPase sites in chromosome condensation. Mol. Cell, 76, 1–14.

Fang, Y. and Spector, D.L. (2005) Centromere positioning and dynamics in living Arabidopsis plants. Mol. Biol. Cell, 16, 5710–5718.

Fazzio, T.G. and Panning, B. (2010) Condensin complexes regulate mitotic progression and interphase chromatin structure in embryonic stem cells. J. Cell Biol., 188, 491–503.

Feng, S., Cokus, S.J., Schubert, V., Zhai, J., Pellegrini, M. and Jacobsen, S.E. (2014) Genome-wide Hi-C analyses in wild-type and mutants reveal high-resolution chromatin interactions in Arabidopsis. Mol. Cell, 55, 694–707.

Fransz, P., de Jong, J.H., Lysak, M., Castiglione, M.R. and Schubert, I. (2002) Interphase chromosomes in Arabidopsis are organized as well defined chromocenters from which euchromatin loops emanate. Proc. Natl. Acad. Sci., 99, 14584–14589.

Freeman, L., Aragon-Alcaide, L. and Strunnikov, A. (2000) The condensin complex governs chromosome condensation and mitotic transmission of rDNA. J. Cell Biol., 149, 811–824.

Fuchs, J., Demidov, D., Houben, A. and Schubert, I. (2006) Chromosomal histone modification patterns – from conservation to diversity. Trends Plant Sci., 11, 199–208.

Fujimoto, S., Yonemura, M., Matsunaga, S., Nakagawa, T., Uchiyama, S. and Fukui, K. (2005) Characterization and dynamic analysis of Arabidopsis condensin subunits, AtCAP-H and AtCAP-H2. Planta, 222, 293–300.

Gentry, M. and Hennig, L. (2014) Remodelling chromatin to shape development of plants. Exp. Cell Res., 321, 40–46.

George, C., Bozler, J., Nguyen, H. and Bosco, G. (2014) Condensins are required for maintenance of nuclear architecture. Cells, 3, 865–882.

Gerlich, D., Hirota, T., Koch, B., Peters, J.M. and Ellenberg, J. (2006) Condensin I stabilizes chromosomes mechanically through a dynamic interaction in live cells. Curr. Biol., 16, 333–344.

Ghavi-Helm, Y., Jankowski, A., Meiers, S., Viales, R., Korbel, J.O. and Furlong, E.M. (2019) Highly rearranged chromosomes reveal uncoupling between genome topology and gene expression. Nature Genetics, 51, 1272–1282.

Gibcus, J.H. and Dekker, J. (2013) The hierarchy of the 3D genome. Mol. Cell, 49, 773–782.

Gibcus, J.H., Samejima, K., Goloborodko, A., Samejima, I., Naumova, N., Nuebler, J., Kanemaki, M.T., Xie, L., Paulson, J.R., Earnshaw, W.C., Mirny, L.A. and Dekker, J. (2018) A pathway for mitotic chromosome formation. Science, 359, 652.

Green, L.C., Kalitsis, P., Chang, T.M., Cipetic, M., Kim, J.H., Marshall, O., Turnbull, L., Whitchurch, C.B., Vagnarelli, P., Samejima, K., Earnshaw, W.C., Choo, K.H. and Hudson, D.F. (2012) Contrasting roles of condensin I and condensin II in mitotic chromosome formation. J. Cell Sci., 125, 1591–1604.

Haase, K., Macadangdang, J.K., Edrington, C.H., Cuerrier, C.M., Hadjiantoniou, S., Harden, J.L., Skerjanc, I.S. and Pelling, A.E. (2016) Extracellular forces cause the nucleus to deform in a highly controlled anisotropic manner. Sci. Rep., 6, 21300.

Habermann, F.A., Cremer, M., Walter, J., Kreth, G., von Hase, J., Bauer, K., Wienberg, J., Cremer, C., Cremer, T. and Solovei, I. (2001) Arrangements of macro- and microchromosomes in chicken cells. Chromosome Res., 9, 569–584.

Hartl, T.A., Sweeney, S.J., Knepler, P.J. and Bosco, G. (2008) Condensin II resolves chromosomal associations to enable anaphase I segregation in Drosophila male meiosis. PLoS Genet., 4, 18–22.

Heckmann, S., Lermontova, I., Berckmans, B., de Veylder, L., Bäumlein, H. and Schubert, I. (2011) The E2F transcription factor family regulates CENH3 expression in *Arabidopsis thaliana*. Plant J., 68, 646–656.

Herzog, S., Jaiswal, S.N., Urban, E., Riemer, A. Fischer, S. and Heidmann, S.K. (2013) Functional dissection of the *Drosophila melanogaster* condensin subunit Cap-G reveals its exclusive association with condensin I. PLoS Genet., 9, e1003463.

Hirano, T. (2012a) Condensins: universal organizers of chromosomes with diverse functions. Genes Dev., 26, 1659–1678.

Hirano, T. (2012b) Chromosome territories meet a condensin. PLoS Genet., 8, e1002939.

Hirota, T., Gerlich, D., Koch, B., Ellenberg, J. and Peters, J.M. (2004) Distinct functions of condensin I and II in mitotic chromosome assembly. J. Cell Sci., 117, 6435–6445.

Hotton, S.K. and Callis, J. (2008) Regulation of cullin RING ligases. Annu. Rev. Plant Biol., 59, 467–489.

Hudson, D.F., Vagnarelli, P., Gassmann, R. and Earnshaw, W.C. (2003) Condensin is required for nonhistone protein assembly and structural integrity of vertebrate mitotic chromosomes. Dev. Cell, 5, 323–336.

Iwasaki, O., Tanaka, A., Tanizawa, H., Grewal, S.I.S. and Noma, K.I. (2010) Centromeric localization of dispersed Pol III genes in fission yeast. Mol. Biol. Cell, 21, 254–265.

Jefferson, R.A., Kavanagh, T.A. and Bevan, M.W. (1987) GUS fusions: B-glucuronidase as a sensitive and versatile gene fusion marker in higher plants. EMBO J., 6, 3901–3907.

Jeppsson, K., Kanno, T., Shirahige, K. and Sjögren, C. (2014) The maintenance of chromosome structure: positioning and functioning of SMC complexes. Nat. Rev. Mol. Cell Biol., 15, 601–614.

Jost, K.L., Bertulat, B. and Cardoso, M.C. (2012) Heterochromatin and gene positioning: inside, outside, any side? Chromosoma, 121, 555–563.

Kagami, Y., Ono, M. and Yoshida, K. (2017) Plk1 phosphorylation of CAP-H2 triggers chromosome condensation by condensin II at the early phase of mitosis. Sci. Rep., 7, 5583.

Kikuchi, S., Kishii, M., Shimizu, M. and Tsujimoto, H. (2005) Centromere-specific repetitive sequences from Torenia, a model plant for interspecific fertilization, and whole-mount FISH of its interspecific hybrid embryos. Cytogenet. Genome Res., 109, 228–235.

Krause, M., Te Riet, J. and Wolf, K. (2013) Probing the compressibility of tumor cell nuclei by combined atomic force-confocal microscopy. Phys. Biol., 10, 065002.

Kudo, T., Sasaki, Y., Terashima, S., Matsuda-Imai, N., Takano, T., Saito, M., Kanno, M., Ozaki, S., Suwabe, K., Suzuki, G., Watanabe, M., Matsuoka, M., Takayama, S. and Yano, K. (2016) Identification of reference genes for quantitative expression analysis using large-scale RNA-seq data of *Arabidopsis thaliana* and model crop plants. Genes Genet. Syst., 91, 111–125.

Lau, A.C., Nabeshima, K. and Csankovszki, G. (2014) The *C. elegans* dosage compensation complex mediates interphase X chromosome compaction. Epigenetics Chromatin, 7, 31.

Layat, E., Sáez-Vásquez, J. and Tourmente, S. (2012) Regulation of pol I-transcribed 45S rDNA and pol III-transcribed 5S rDNA in Arabidopsis. Plant Cell Physiol., 53, 267–276.

Liu, C., Wang, C., Wang, G., Becker, C., Zaidem, M. and Weigel, D. (2016) Genome-wide analysis of chromatin packing in *Arabidopsis thaliana* at single-gene resolution. Genome Res., 26, 1057–1068.

Liu, C.M., McElver, J., Tzafrir, I., Joosen, R., Wittich, P., Patton, D., van Lammeren, A.A.M. and Meinke, D. (2002) Condensin and cohesin knockouts in Arabidopsis exhibit a titan seed phenotype. Plant J., 29, 405–415.

Livak, K.J. and Schmittgen, T.D. (2001) Analysis of relative gene expression data using real-time quantitative PCR and the 2-^ΔΔC^T method. Methods, 25, 402–408.

Longworth, M.S. Herr, A., Ji, J.Y. and Dyson, N.J. (2008) RBF1 promotes chromatin condensation through a conserved interaction with the Condensin II protein dCAP-D3. Genes Dev., 22, 1011–1024.

Longworth, M.S., Walker, J.A., Anderssen, E., Moon, N.S., Gladden, A., Heck, M.M.S., Ramaswamy, S. and Dyson, N.J. (2012) A shared role for RBF1 and dCAP-D3 in the regulation of transcription with consequences for innate immunity. PLoS Genet., 8, e1002618.

Love, M.I., Huber, W. and Anders, S. (2014) Moderated estimation of fold change and dispersion for RNA-seq data with DESeq2. Genome Biol., 15, 550.

Maeshima, K. and Laemmli, U.K. (2003) A two-step scaffolding model for mitotic chromosome assembly. Dev Cell 4: 467–480.

Maeshima, K., Tamura, S. and Shimamoto, Y. (2018) Chromatin as a nuclear spring. Biophys. Physicobiol., 15, 189–195.

Marenduzzo, D., Finan, K. and Cook, P.R. (2006) The depletion attraction: An underappreciated force driving cellular organization. J. Cell Biol., 175, 681–686.

Martinez-Zapater, J.M., Estelle, M.A. and Somerville, C.R. (1986) A highly repeated DNA sequence in Arabidopsis thaliana. Mol. Gen. Genet., 204, 417–423.

Mayer, R., Brero, A., von Hase, J., Schroeder, T., Cremer, T. and Dietzel, S. (2005) Common themes and cell type specific variations of higher order chromatin arrangements in the mouse. BMC Cell Biol., 6, 44.

Moissiard, G., Cokus, S.J., Cary, J., Feng, S., Billi, A.C., Stroud, H., Husmann, D., Zhan, Y., Lajoie, B.R., McCord, R.P., Hale, C.J., Feng, W., Michaels, S.D., Frand, A.R., Pellegrini, M., Dekker, J., Kim, J.K. and Jacobsen, S.E. (2012) MORC family ATPases required for heterochromatin condensation and gene silencing. Science, 336, 1448–1451.

Nakamura, S., Mano, S., Tanaka, Y., Ohnishi, M., Nakamori, C., Araki, M., Niwa, T., Nishimura, M., Kaminaka, H., Nakagawa, T., Sato, Y. and Ishiguro, S. (2010) Gateway binary vectors with the bialaphos resistance gene, bar, as a selection marker for plant transformation. Biosci. Biotechnol. Biochem., 74, 1315–1319.

Onn, I., Aono, N., Hirano, M. and Hirano, T. (2007) Reconstitution and subunit geometry of human condensin complexes. EMBO J., 26, 1024–1034.

Ono, T., Fang, Y., Spector, D.L. and Hirano, T. (2004) Spatial and temporal regulation of condensins I and II in mitotic chromosome assembly in human cells. Mol. Biol. Cell, 15, 3296–3308.

Ono, T., Losada, A., Hirano, M., Myers, M.P., Neuwald, A.F. and Hirano, T. (2003) Differential contributions of condensin I and condensin II to mitotic chromosome architecture in vertebrate cells. Cell, 115, 109–121.

Parra, G., Bradnam, K., Rose, A.B. and Korf, I. (2011) Comparative and functional analysis of intron-mediated enhancement signals reveals conserved features among plants. Nucleic Acids Res., 39, 5328–5337.

Pecinka, A., Schubert, V., Meister, A., Kreth, G., Klatte, M., Lysak, M.A., Fuchs, J. and Schubert, I. (2004) Chromosome territory arrangement and homologous pairing in nuclei of *Arabidopsis thaliana* are predominantly random except for NOR-bearing chromosomes. Chromosoma, 113, 258–269.

Perrella, G., Lopez-Vernaza, M.A., Carr, C., Sani, E., Gossele, V., Verduyn, C., Kellermeier, F., Hannah, M.A. and Amtmann, A. (2013) Histone deacetylase complex1 expression level titrates plant growth and abscisic acid sensitivity in Arabidopsis. Plant Cell, 25, 3491–3505.

Poulet, A., Duc, C., Voisin, M., Desset, S., Tutois, S., Vanrobays, E., Benoit, M., Evans, D.E., Probst, A.V. and Tatout, C. (2017) The LINC complex contributes to heterochromatin organisation and transcriptional gene silencing in plants. J. Cell Sci., 130, 590–601.

Robellet, X., Fauque, L., Legros, P., Mollereau, E., Janczarski, S., Parrinello, H., Desvignes, J.P., Thevenin, M. and Bernard, P. (2014) A genetic screen for functional partners of condensin in fission yeast. G3 Genes|Genomes|Genetics, 4, 373–381.

Robson M.I., Ringel A.R., and Mundlos S. (2019) Regulatory landscaping: how enhancer-promoter communication is sculpted in 3D. Mol. Cell, 74, 1110–1122.

Rose, A.B., Elfersi, T., Parra, G. and Korf, I. (2008) Promoter-proximal introns in *Arabidopsis thaliana* are enriched in dispersed signals that elevate gene expression. Plant Cell, 20, 543–551.

Rosin, L.F., Nguyen, S.C. and Joyce, E.F. (2018) Condensin II drives large-scale folding and spatial partitioning of interphase chromosomes in Drosophila nuclei. PLoS Genet., 14, e1007393.

Probst, A.V. and Mittelsten-Scheid, O. (2015) Stress-induced structural changes in plant chromatin. Curr. Opin. Plant. Biol., 27, 8–16.

Sakamoto, T., Sugiyama, T., Yamashita, T. and Matsunaga, S. (2019) Plant condensin II is required for the correct spatial relationship between centromeres and rDNA arrays. Nucleus., 10, 116–125.

Sakamoto, Y. and Takagi, S. (2013) LITTLE NUCLEI 1 and 4 regulate nuclear morphology in *Arabidopsis thaliana*. Plant Cell Physiol., 54, 622–633.

Savvidou, E., Cobbe, N., Steffensen, S., Cotterill, S. and Heck, M.M.S. (2005) Drosophila CAP-D2 is required for condensin complex stability and resolution of sister chromatids. J. Cell Sci., 118, 2529–2543.

Schägger, H. and von Jagow, G. (1987) Tricine-sodium dodecyl sulfate-polyacrylamide gel electrophoresis for the separation of proteins in the range from 1 to 100 kDa. Anal. Biochem., 166, 368–379.

Schmiesing, J., Gregson, H., Zhou, S. and Yokomori, K. (2000) A human condensin complex containing hCAP-C-hCAP-E and CNAP1, a homolog of Xenopus XCAP-D2, colocalizes with phosphorylated histone H3 during the early stage of mitotic chromosome condensation. Mol. Cell. Biol., 20, 6996–7006.

Schneider, C.A., Rasband, W.S. and Eliceiri, K.W. (2012) NIH Image to ImageJ: 25 years of image analysis. Nat. Meth., 9, 671–675.

Schubert, V. (2009) SMC proteins and their multiple functions in higher plants. Cytogenet. Genome Res., 124, 202–214.

Schubert, V., Berr, A. and Meister, A. (2012) Interphase chromatin organisation in Arabidopsis nuclei: constraints versus randomness. Chromosoma, 121, 369–387.

Schubert, V., Lermontova, I. and Schubert, I. (2013) The Arabidopsis CAP-D proteins are required for correct chromatin organisation, growth and fertility. Chromosoma, 122, 517–533.

Shao, Y., Lu, N., Wu, Z., Cai, C., Wang, S., Zhang, L., Zhou, F., Xiao, S., Liu, L., Zeng, X., Zheng, H., Yang, C., Zhao, Z., Zhao, G., Zhou, J., Xue, X., Qin, Z. (2018) Creating a functional single-chromosome yeast. Nature, 560, 331–335.

Siddiqui, N.U., Stronghill, P.E., Dengler, R.E., Hasenkampf, A. and Riggs, C.D. (2003) Mutations in Arabidopsis condensin genes disrupt embryogenesis, meristem organization and segregation of homologous chromosomes during meiosis. Development, 130, 3283–3295.

Sieburth, L.E. and Meyerowitz, E.M. (1997) Molecular dissection of the AGAMOUS control region shows that cis elements for spatial regulation are located intragenically. Plant Cell, 9, 355–365.

Skibbens, R.V. (2019) Condensins and cohesins - one of these things is not like the other! J. Cell Sci., 132, jcs220491.

Skylar, A., Matsuwaka, S. and Wu, X. (2013) ELONGATA3 is required for shoot meristem cell cycle progression in *Arabidopsis thaliana* seedlings. Dev. Biol., 382, 436–445.

Smith, S.J., Osman, K. and Franklin, F.C.H. (2014) The condensin complexes play distinct roles to ensure normal chromosome morphogenesis during meiotic division in Arabidopsis. Plant J., 80, 255–268.

Soppe, W.J.J., Jasencakova, Z., Houben, A., Kakutani, T., Meister, A., Huang, M.S., Jacobsen, S.E., Schubert, I. and Fransz, P.F. (2002) DNA methylation controls histone H3 lysine 9 methylation and heterochromatin assembly in Arabidopsis. EMBO J., 21, 6549–6559.

Sparkes, I.A., Runions, J., Kearns, A. and Hawes, C. (2006) Rapid, transient expression of fluorescent fusion proteins in tobacco plants and generation of stably transformed plants. Nat. Protoc., 1, 2019–2025.

Stam M., Tark-Dame M. and Fransz P. (2019) 3D genome organization: a role for phase separation and loop extrusion? Curr. Opin. Plant Biol., 48, 36–46.

Sun, M., Biggs, R., Hornick, J. and J.F. Marko (2018) Condensin controls mitotic chromosome stiffness and stability without forming a structurally contiguous scaffold. Chromosome Res., 26, 277–295.

Szabo, Q., Bantignies, F. and Cavalli G. (2019) Principles of genome folding into topologically associating domains. Sci. Adv, 5, eaaw1668.

Szklarczyk, D., Gable, A.L., Lyon, D., Junge, A., Wyder, S., Huerta-Cepas, J., Simonovic, M., Doncheva, N.T., Morris, J.H., Bork, P., Jensen, L.J. and Mering, C.V. (2019) STRING v11: protein-protein association networks with increased coverage, supporting functional discovery in genome-wide experimental datasets. Nucleic Acids Res., 47, D607–D613.

Tatout, C., Evans, D.E., Vanrobays, E., Probst, A.V. and Graumann, K. (2014) The plant LINC complex at the nuclear envelope. Chromosom. Res., 22, 241–252.

Tessadori, F., Chupeau, M.C., Chupeau, Y., Knip, M., Germann, S., van Driel, R., Fransz, P. and Gaudin, V. (2007) Large-scale dissociation and sequential reassembly of pericentric heterochromatin in dedifferentiated Arabidopsis cells. J. Cell Sci., 120, 1200–1208.

Van Leene, J., Eeckhout, D., Cannoot, B., De Winne, N., Persiau, G., Van De Slijke, E., Vercruysse, L., Dedecker, M., Verkest, A., Vandepoele, K., Martens, L., Witters, E., Gevaert, K. and De Jaeger, G. (2014) An improved toolbox to unravel the plant cellular machinery by tandem affinity purification of Arabidopsis protein complexes. Nat. Protoc., 10, 169–187.

Van Leene, J., Eeckhout, D., Persiau, G., van de Slijke, E., Geerinck, J., van Isterdael, G., Witters, E. and de Jaeger, G. (2011) Isolation of transcription factor complexes from Arabidopsis cell suspension cultures by tandem affinity purification. In L. Yuan and S. E. Perry, eds. Plant transcription factors. Meth. Mol. Biol., 195–218.

Van Ruiten, M.S. and Rowland, B.D. (2018) SMC Complexes: universal DNA looping machines with distinct regulators. Trends Genet., 34, 477–487.

Van Zanten, M., Koini, M.A., Geyer, R., Liu, Y., Brambilla, V., Bartels, D., Koornneef, M., Fransz, P., and Soppe W.J. (2011) Seed maturation in *Arabidopsis thaliana* is characterized by nuclear size reduction and increased chromatin condensation. Proc. Natl. Acad. Sci. U S A, 108, 20219–20224.

Verlinden, L., Eelen, G., Beullens, I., Van Camp, M., Van Hummelen, P., Engelen, K., Van Hellemont, R., Marchal, K., De Moor, B., Foijer, F., Te Riele, H., Beullens, M., Bollen, M., Mathieu, C., Bouillon, R. and Verstuyf, A. (2005) Characterization of the condensin component Cnap1 and protein kinase melk as novel E2F target genes down-regulated by 1,25-dihydroxyvitamin D3. J. Biol. Chem., 280, 37319–37330.

Wallace, H.A. and Bosco, G. (2013) Condensins and 3D organization of the interphase nucleus. Curr. Genet. Med. Rep., 1, 219–229.

Wallace, H.A., Klebba, J.E., Kusch, T., Rogers, G.C. and Bosco, G. (2015) Condensin II regulates interphase chromatin organization through the Mrg-binding motif of Cap-H2. G3 Genes/Genomes/Genetics 5, 803–817.

Walther, N., Hossain, M.J., Politi, A.Z., Koch, B., Kueblbeck, M., Odegard-Fougner, O., Lampe, M. and Ellenberg, J. (2018) A quantitative map of human condensins provides new insights into mitotic chromosome architecture. J. Cell Biol., 217, 2309–2328.

Wang, H., Dittmer, T.A. and Richards, E.J. (2013) Arabidopsis CROWDED NUCLEI (CRWN) proteins are required for nuclear size control and heterochromatin organization. BMC Plant Biol., 13, 200.

Wang, J., Blevins, T., Podicheti, R., Haag, J.R., Tan, E.H., Wang, F. and Pikaard, C.S. (2017) Mutation of Arabidopsis SMC4 identifies condensin as a corepressor of pericentromeric transposons and conditionally expressed genes. Genes Dev., 31, 1601–1614.

Wang, Z., Cao, H., Chen, F. and Liu, Y. (2014) The roles of histone acetylation in seed performance and plant development. Plant Physiol. Biochem., 84, 125–133.

Weisshart, K., Fuchs, J. and Schubert, V. (2016) Structured illumination microscopy (SIM) and photoactivated localization microscopy (PALM) to analyze the abundance and distribution of RNA polymerase II molecules on flow-sorted Arabidopsis nuclei. Bio-Protocol, 6. e1725.

Winter, D., Vinegar, B., Nahal, H., Ammar, R., Wilson, G.V. and Provart, N.J. (2007) An “Electronic Fluorescent Pictograph” browser for exploring and analyzing large-scale biological data sets. PLoS One, 8, e718.

Wu, T.D. and Nacu, S. (2010) Fast and SNP-tolerant detection of complex variants and splicing in short reads. Bioinformatics, 26, 873–881.

Xia, Y., Pfeifer, C.R., Cho, S., Discher, D.E. and Irianto, J. (2018) Nuclear mechanosensing. Emerg. Top. Life Sci., 2, 713–725.

Yilmaz, A., Mejia-Guerra, M.K., Kurz, K., Liang, X., Welch, L. and Grotewold E. (2011) AGRIS: the Arabidopsis gene regulatory information server, an update. Nucleic Acids Res., 39, D1118–D1122.

Yoo, S.D., Cho, Y.H. and Sheen, J. (2007) Arabidopsis mesophyll protoplasts: a versatile cell system for transient gene expression analysis. Nat. Protoc., 2, 1565–1572.

Yuen, K.C., Slaughter, B.D. and Gerton, J.L. (2017) Condensin II is anchored by TFIIIC and H3K4me3 in the mammalian genome and supports the expression of active dense gene clusters. Sci. Adv., 3, e1700191.

Zamariola, L., de Storme, N., Vannerum, K., Vandepoele, K., Armstrong, S.J., Franklin, F.C.H. and Geelen, D. (2014) SHUGOSHINs and PATRONUS protect meiotic centromere cohesion in *Arabidopsis thaliana*. Plant J., 77, 782–794.

Zelkowski, M., Zelkowska, K., Conrad, U., Hesse, S., Lermontova, I., Marzec, M., Meister, A., Houben, A. and Schubert, V. (2019) *Arabidopsis* NSE4 proteins act in somatic nuclei and meiosis to ensure plant viability and fertility. Front. Plant Sci., 10, 774.

Zhang, T., Paulson, J.R., Bakhrebah, M., Kim, J.H., Nowell, C., Kalitsis, P. and Hudson, D.F. (2016) Condensin I and II behaviour in interphase nuclei and cells undergoing premature chromosome condensation. Chromosom. Res., 24, 243–269.

